# A leptin-responsive hypothalamic circuit inputs to the circadian feeding network

**DOI:** 10.1101/2023.02.24.529901

**Authors:** Qijun Tang, Elizabeth Godschall, Charles D. Brennan, Qi Zhang, Ruei-Jen Abraham-Fan, Sydney P. Williams, Taha Buğra Güngül, Roberta Onoharigho, Aleyna Buyukaksakal, Ricardo Salinas, Joey J. Olivieri, Christopher D. Deppmann, John N. Campbell, Brandon Podyma, Ali D. Güler

**Author notes:** Equal contribution (Qijun Tang, Elizabeth Godschall, Brandon Podyma). <u>Correspondence</u> To whom correspondence should be sent: Ali D. Güler, Departments of Biology and Neuroscience University of Virginia, Charlottesville, VA 22904; Brandon Podyma, Medical Scientist Training Program, School of Medicine University of Virginia, Charlottesville, VA 22903; John N. Campbell, Departments of Biology and Neuroscience University of Virginia, Charlottesville, VA 22904.

## Abstract

Salient cues, such as the rising sun or the availability of food, play a crucial role in entraining biological clocks, allowing for effective behavioral adaptation and ultimately, survival. While the light-dependent entrainment of the central circadian pacemaker (suprachiasmatic nucleus, SCN) is relatively well defined, the molecular and neural mechanisms underlying entrainment associated with food availability remains elusive. Using single nucleus RNA sequencing during scheduled feeding (SF), we identified a leptin receptor (LepR) expressing neuron population in the dorsomedial hypothalamus (DMH) that upregulates circadian entrainment genes and exhibits rhythmic calcium activity prior to an anticipated meal. We found that disrupting DMH^LepR^ neuron activity had a profound impact on both molecular and behavioral food entrainment. Specifically, silencing DMH^LepR^ neurons, mis-timed exogenous leptin administration, or mis-timed chemogenetic stimulation of these neurons all interfered with the development of food entrainment. In a state of energy abundance, repetitive activation of DMH^LepR^ neurons led to the partitioning of a secondary bout of circadian locomotor activity that was in phase with the stimulation and dependent on an intact SCN. Lastly, we discovered that a subpopulation of DMH^LepR^ neurons project to the SCN with the capacity to influence the phase of the circadian clock. This leptin regulated circuit serves as a point of integration between the metabolic and circadian systems, facilitating the anticipation of meal times.

## Introduction

When we eat is as important for our health as what and how much we eat. Studies in both mice and humans have shown that eating during the rest phase (daytime for mice or nighttime for humans) is associated with increased risk of weight gain, glucose intolerance, hepatic steatosis, and cardiovascular disease ^1–4^. Efforts to mitigate these deleterious effects in mice by restricting when they eat have provided significant promise to improve metabolic health and even extend lifespan. The potential benefits of time-restricted eating to human cardiometabolic health is the focus of many ongoing clinical studies ^5–9^. However, we have a limited mechanistic understanding of how meal timing influences our physiology and biological rhythms ^10^. Therefore, we sought to better understand the anatomical and molecular underpinnings of the interaction between feeding time and the circadian clock using a model of scheduled feeding (SF) that rapidly induces biological entrainment in rodents ^11–13^.

The suprachiasmatic nucleus (SCN) in the hypothalamus is the primary pacemaker that receives ambient light information from the retina, synchronizes circadian machinery throughout the body, and coordinates behavioral outputs ^14,15^. Interestingly and less well understood, in the absence of a functional SCN ^12,13^, the circadian system retains the ability to entrain to the timing of non-photic environmental cues, such as food ^11^. Numerous efforts have failed to identify any necessary genetic, molecular, or anatomic substrates of food entrainment ^16,17^. Emerging evidence suggests that the food entrainment system encompasses multiple food entrainable oscillators distributed across the central nervous system and peripheral organs, in which partial malfunction is compensated for by other parts of the network (Fig 1A) ^10,17–21^.

**Figure 1.**
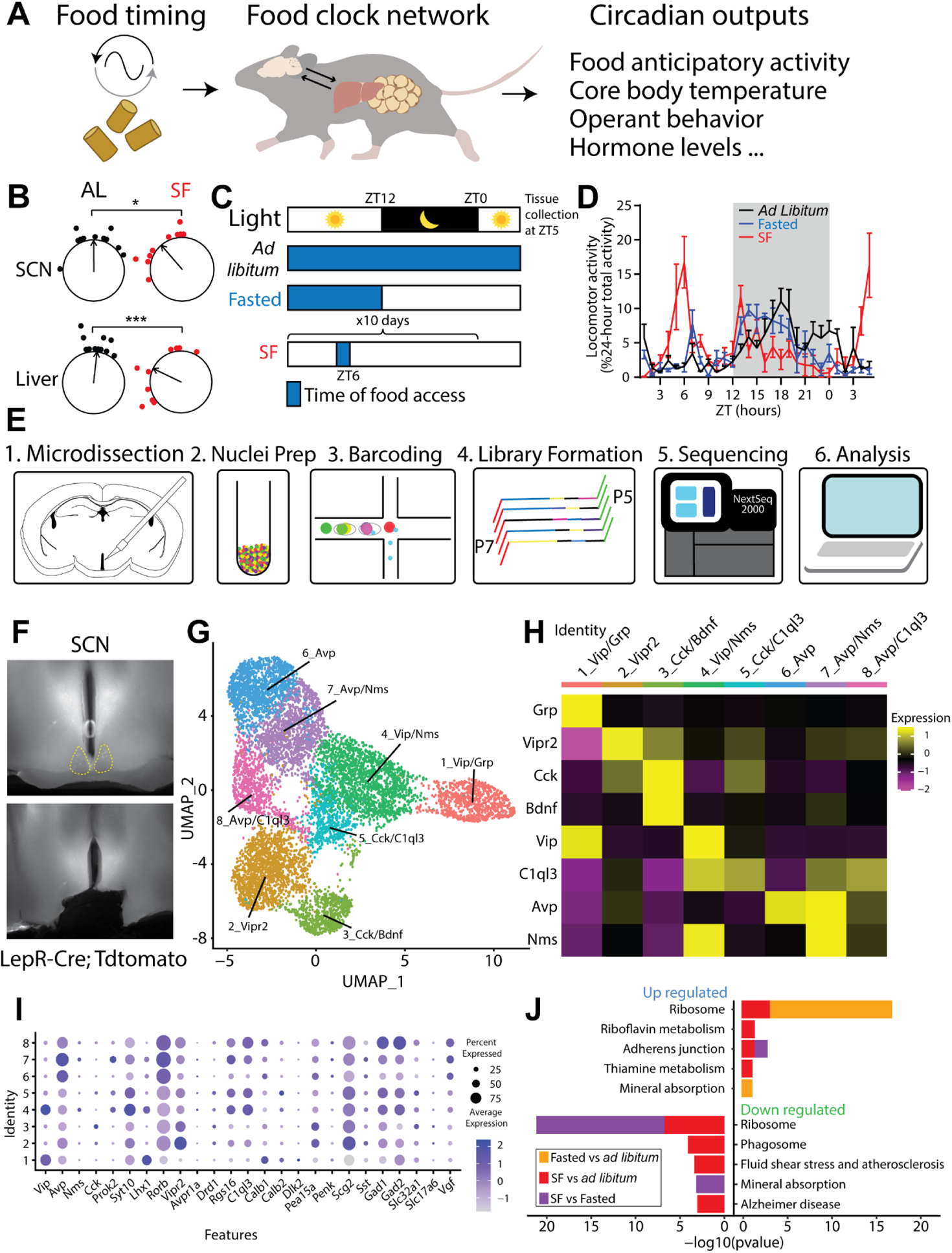
SCN snRNAseq reveals minimal alteration of circadian genes during SF. **A.** Diagram illustrating that food timing as a potent zeitgeber entraining an oscillatory network system in the brain and peripheral organs, relaying rhythmic behavior outputs. **B.** The ZT phase of the first bioluminescence peak of SCN and liver from PER2::Luciferase mice that are either provided with scheduled food access for 4 days at ZT6 or *ad libitum* fed controls (untreated or given ZT6 saline injections). Two-way ANOVA with Bonferroni post hoc comparison; n = 10-11 / group; F_treatment_ (1, 38) = 19.05, p<0.001. **C.** Schematic of experimental design. Mice were housed on a 12-12 light-dark cycle and either fed *ad libitum,* overnight fasted, or provided a scheduled meal for 10 days at ZT6. Blue shading denotes food access. All mice were sacrificed for tissue collection at ZT5. **D.** Normalized locomotor activity starting 29 hours before tissue collection. n=4 mice / condition. Data are represented as mean ± SEM. **E.** Schematic of single nuclei RNA sequencing (snRNAseq) workflow using 10X Genomics. **F.** Representative images illustrating the area of dissection in the SCN for snRNAseq. **G.** Uniform Manifold Approximation and Projection (UMAP) plot of 8 molecularly distinct SCN neuron subtypes (n=8,957 neurons). **H.** Heatmap of cluster-average marker gene expression, scaled by gene. **I.** Dot plot of average expression level (dot color) and percent expression (dot size) for each SCN neuron cluster. Genes shown were previously defined as SCN markers ^35,36^ and validated based on Allen Brain Atlas Mouse Brain *in situ* hybridization data ^37^. **J.** Kyoto Encyclopedia of Genes and Genomes (KEGG) from the Mouse 2019 database comparing top 5 pathways up- and down-regulated among feeding conditions in all SCN neurons. Inclusion criteria required p-value <0.05 and log2 fold change >0.25. See also supplemental figure 1.

Here we used a time and calorie restricted feeding paradigm to rapidly induce food entrainment in mice ^22^, with a focus on the SCN and the dorsomedial hypothalamus (DMH) which are involved in the regulation of feeding, locomotor activity, sleep-wake cycles, and hormone rhythms ^23–26^. By using single nucleus RNA sequencing, we sought to identify neuronal populations in the SCN and DMH that show changes in circadian transcriptional programs. We did not observe appreciable transcriptional changes of genes associated with circadian rhythmicity or circadian entrainment pathways in the SCN during scheduled feeding (SF). However, we identified several neuronal populations in the DMH that altered their expression of circadian entrainment genes in response to timed feeding, including the leptin-receptor (LepR) expressing neurons. Next, we demonstrated that chronic silencing or mis-timed over-activation of the DMH^LepR^ neurons, as well as mis-timed leptin administration, impair development of food entrainment. Finally, we uncovered a direct neuronal projection from DMH^LepR^ neurons to the SCN and showed that DMH^LepR^ neuron stimulation is sufficient to phase shift the SCN circadian clock while altering the structure of circadian locomotor activity. These results define a mechanism that integrates mealtime information with the circadian clock via leptin signaling in the DMH.

## Results

### Scheduled feeding alters “circadian entrainment” gene expression in the DMH but not the SCN

In mammals, the SCN in the hypothalamus is the seat of the primary circadian clock which receives ambient light signals and synchronizes the biological clocks distributed throughout the body ^14^. Although it is not required for the expression of food anticipatory behavior (FAA) ^13,27,28^, the SCN has recently been shown to modify the robustness of food entrainment (measured by one of the behavioral outputs of food entrainment, FAA) ^29^ (Fig 1A). We tested the entrainment of central and peripheral circadian systems using a scheduled feeding (SF) paradigm where we restricted both time and calories of food delivered. In contrast to only time-restricted feeding regimens, which shift peripheral but not central circadian clocks ^30^, this SF paradigm induces a phase advance in the bioluminescent reported circadian rhythmicity of both the SCN and the liver from the PER2::Luciferase (PER2LUC) transgenic mice (Fig 1B) ^31^. Our observation is in line with previously demonstrated SCN rhythm phase shifts in time- and calorie-restricted animals ^32,33^. To further elucidate the transcriptional programs of hypothalamic regions in food entrainment, we harvested fresh brain tissues at 5 hours after lights on (*Zeitgeber* time or ZT 5) from mice that were subjected to three feeding conditions: *ad libitum*, overnight fasted, or fed at ZT 6 for ten days (SF; Fig 1C-D). We isolated SCN, as well as DMH, a hypothalamic region previously implicated in circadian and feeding regulation (Fig 1E) ^23,34^. After brain region- and feeding condition-specific tissue collection and nuclei isolation, we performed single-nucleus RNA sequencing (snRNAseq), yielding raw datasets of 59,708 and 65,837 cells from SCN- and DMH-containing tissues, respectively.

Using previously defined SCN markers (e.g., *Avp, Vip, Vipr2, Prok2, Per2)* ^35,36^, we identified 8,957 cells as SCN neurons and clustered them by transcriptomic similarity into 8 candidate subtypes (Fig 1F-I, Supplemental Fig 1A-D). In the final SCN dataset, the mean number of genes and unique transcripts (unique molecular identifiers, UMIs) detected per cell in all SCN samples was 1,783 and 3,294, respectively (Supplemental Fig 1C). We then compared SCN neuron gene expression across feeding conditions: SF versus *ad libitum*, SF versus fasting, and fasting versus *ad libitum* conditions. Using the Kyoto Encyclopedia of Genes and Genomes (KEGG) Mouse 2019 database, we identified the top five up- and down-regulated pathways in the SCN (Fig 1J). Neither circadian entrainment nor circadian rhythm pathways were significantly altered in the SCN under SF relative to the other feeding conditions. Despite the primacy of the SCN in photic based pacemaking and the shift observed in Per2 rhythmicity (Fig 1B), these snRNAseq data show that transcriptional alteration of the circadian system is limited in the SCN in response to food based pacemaking, in line with previous work showing its expendability for food entrainment ^12,13,27^.

Using previously defined SCN markers (e.g. *Avp, Vip, Vipr2, Prok2, Per2)* ^35,36^, 8,957 cells were identified as SCN neurons and grouped into 8 clusters (Fig 1F-I, Supplemental Fig 1A-D). In the final SCN dataset, the mean number of genes and unique molecular identifiers (UMIs) detected per cell in SCN samples was 1,783 and 3,294, respectively (Supplemental Fig 1C). We then compared SCN neurons of SF versus *ad libitum*, SF versus fasting, and fasting versus *ad libitum* conditions via differential expression analysis. Using the Kyoto Encyclopedia of Genes and Genomes (KEGG) Mouse 2019 database, we identified the top five up- and down-regulated pathways in the SCN (Fig 1J). We did not identify mediators of circadian entrainment or circadian rhythms as significantly altered in the SCN under SF. Despite the primacy of the SCN in photic based pacemaking and the shift observed in Per2 rhythmicity (Fig 1B), these snRNAseq data show that transcriptional alteration of the circadian system is limited in the SCN in response to food based pacemaking, in line with previous work showing its expendability for food entrainment^12,13,27^.

We next turned our attention to the DMH, a neighboring hypothalamic region which has been strongly implicated in circadian behaviors and physiological processes ^23–26^. However, the extent of DMH involvement in food entrainment is controversial with substantial reproducibility concerns ^17,18,26, 38–50^, potentially due to the heterogeneity of DMH neurons. Therefore, we used snRNAseq to compare the gene expression profiles of 16,281 DMH neurons from mice under *ad libitum*, fasted, or scheduled feeding conditions. We first identified DMH neurons from our snRNA-seq dataset based on their enriched expression of known DMH markers including *Gpr50, Grp, Rorb, Sulf1, Pcsk5, Lepr, Pdyn,* and *Ppp1r17* ^45,51,52^. We then clustered these putative DMH neurons into 14 candidate subtypes according to transcriptomic similarity and annotated them based on top marker genes (Fig 2A-D, Supplemental Fig 1E-H). The mean number of genes and UMIs per cell detected in all DMH samples was 2,425 and 5,235, respectively (Supplemental Fig 1G). Our dataset contained clusters corresponding to previously identified DMH neuron populations, those expressing *Lepr*, *Pdyn*, or *Ppp1r17*^43–45,52^, along with novel cluster-specific expression of 2_Tcf7l2 ^53^ or 12_Nfix that were together named *Lhx6*+ neurons previously^54^. In sharp contrast to the SCN, DMH KEGG pathway analysis revealed upregulation of the “circadian entrainment” genes (e.g. *Kcnj6, Gria2, Nos1*, etc.), which are involved in transmitting salient extracellular signaling cues to the core molecular clock (https://www.kegg.jp/entry/map04713; Supplemental Fig 1I). This upregulation was seen not only in SF vs *ad libitum*, but also SF vs fasted conditions, demonstrating that the effect on expression of circadian entrainment genes was not simply due to energy deficit, but adaptation to food timing (Fig 2E). These results imply that the genes capable of influencing the DMH circadian clock are altered by scheduled feeding.

**Figure 2.**
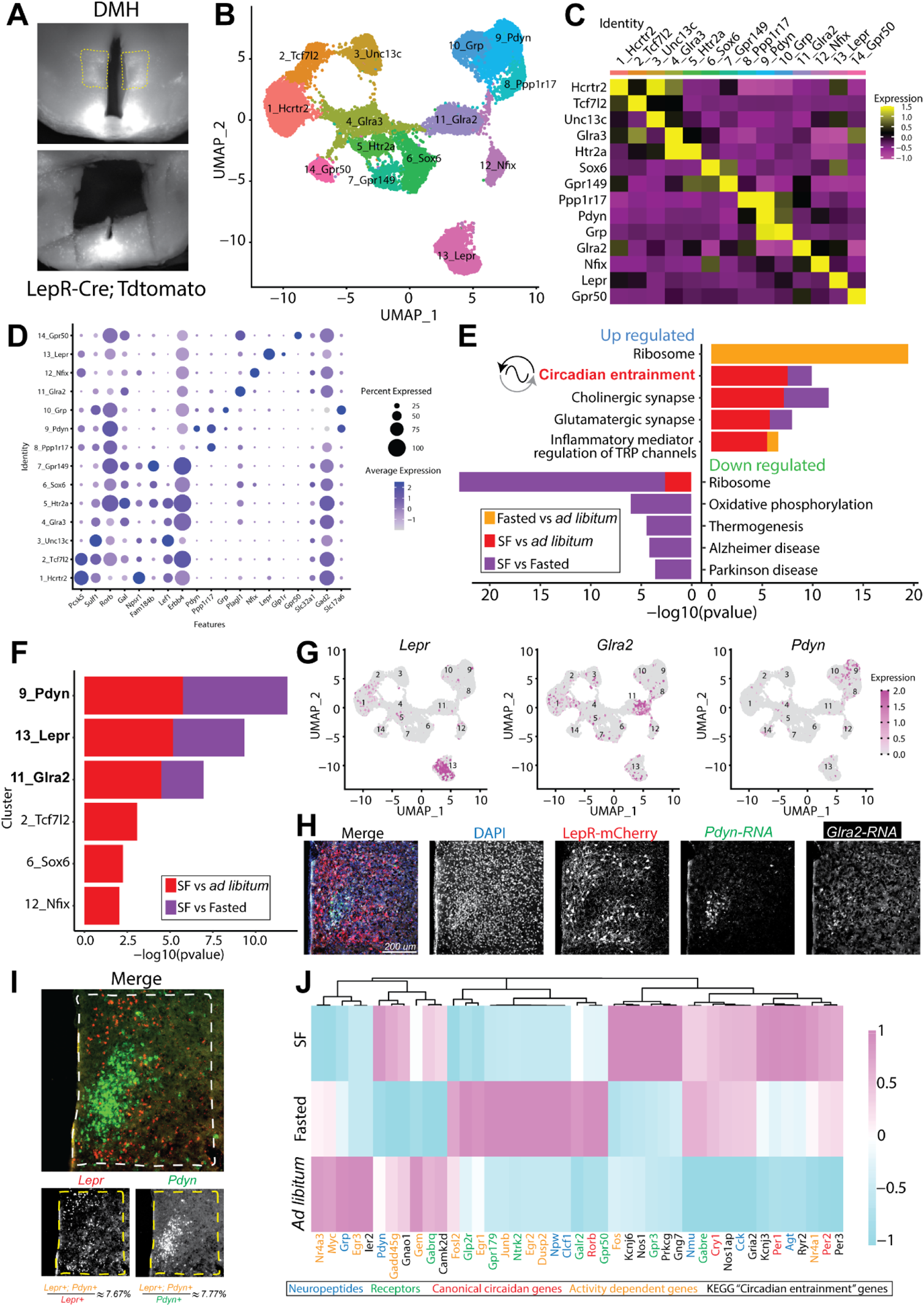
SF alters circadian entrainment genes in specific DMH neuron subtypes. **A.** Representative images illustrating the DMH area dissected for snRNAseq. **B.** UMAP of 14 defined DMH neuron subtypes (n=16,281 neurons). **C.** Average gene expression heatmap labeled by cluster-specific markers in the DMH. **D.** Dot plot of average expression level (dot color) and percent expression (dot size) of genes of interest within DMH clusters. These genes were either previously identified in DMH ^45,51,52,57^ or validated as DMH markers by the Allen Brain Atlas Mouse Brain *in situ* hybridization data^37^. **E.** KEGG from the Mouse 2019 database comparing top 5 pathways up- and down-regulated across feeding conditions in all DMH neurons. Inclusion criteria required p-value <0.05 and log2 fold change >0.25. **F.** DMH clusters with differentially regulated circadian entrainment pathways in at least one scheduled feeding comparison. **G.** Feature plots indicating spatial expression of *Lepr* (left), *Glra2* (middle), *Pdyn* (right), in DMH clusters. **H.** Representative coronal section image localizing expression of LepR, *Pdyn*, and *Glra2* in the DMH. LepR cells were marked by LepR-Cre;TdTomato protein, whereas *Pdyn* and *Glra2* transcripts were visualized by RNA FISH. See also supplemental Fig 3G for zoomed-out view of the same brain section. **I.** Representative RNA FISH coronal section image showing *Lepr* and *Pdyn* transcripts in the DMH. Quantification of *Lepr* and *Pdyn* co-expressing cells is depicted at the bottom. n=3 mice. **J.** Heatmap of select genes that were differentially expressed across feeding conditions in DMH^LepR^ neurons. See also supplemental figure 1 and 2.

### SF alters circadian entrainment gene expression in DMH^LepR^ neurons

The DMH is a heterogeneous and ill-defined anatomic area containing numerous genetically distinct cell populations, two of which (expressing either *Pdyn* or *Ppp1r17*) have been previously investigated in food entrainment behavior ^43–45^. Thus, we sought to understand which DMH neuronal subpopulations exhibit the most significant change in “circadian entrainment” gene expression during SF. Of the 14 neuron clusters we identified in the DMH, six showed differential gene expression in circadian entrainment pathway during energy deficit, and three of these had differential gene expression in both SF vs. *ad libitum* and SF vs. fasted conditions: cluster 9, Pdyn [prodynorphin] neurons; cluster 13, Lepr [leptin receptor] neurons; and cluster 11, Glra2 [glycine receptor subunit alpha-2] neurons (Fig 2F-H, Supplemental Fig 2A).

Of these candidate DMH neuron subtypes, those expressing *Pdyn* and *Lepr* have putative connections with both circadian and feeding regulation ^24,43^, and exhibit strikingly different anatomic distributions within the DMH, while *Glra2* does not (Fig 2H, Supplemental Fig 2E-G). For these reasons we chose to further investigate the *Pdyn+* and *Lepr+* neuron subtypes in our dataset. *Lepr+* and *Pdyn+* neurons partially overlap in the DMH ^52^. However, using RNA fluorescence *in situ* hybridization (RNA FISH), we found that this overlap is minimal: only ~7.67% of *Lepr+* neurons are *Pdyn*+, while ~7.77% of *Pdyn+* neurons are *Lepr*+ (Fig 2I). Additionally, we observed that the *Lepr* expression is predominant in the dorsal and ventral DMH, whereas *Pdyn* expression is confined to the compact central DMH (Fig 2I). The isolated “core/shell” expression pattern of the two populations of neurons suggests that they play distinct functional roles in food entrainment in the DMH. The DMH^Pdyn^ neurons have been shown to entrain to scheduled feeding^43^ and dampen the robustness of FAA when silenced ^44^. As one of the major sources of inhibitory input to the agouti-related peptide (AgRP) neurons of the arcuate nucleus, DMH^LepR^ neurons are important for feeding and energy homeostasis ^24,52,55,56^. However, their contribution to food entrainment is unknown. Detailed analysis of the DMH cluster 13 Lepr neurons in our dataset revealed that SF alters transcription of circadian entrainment pathway genes (Supplemental Fig 1I, 2D), as well as activity dependent genes, neuropeptides, receptors, and canonical circadian genes. Together these transcriptional responses to scheduled feeding point to a significant role for DMH^LepR^ neurons in food entrainment ^43,44^ (Fig 2J, Supplemental Fig 2A-D).

### Leptin suppresses food anticipatory behavior- and the calcium-activity of DMH^LepR^ neurons

Since leptin is the ligand for LepR, we next sought to examine the effect of leptin on food entrainment. Predominantly made by adipose tissue, leptin is released in response to a meal ^58^ and scaled to the time of day ^59–61^. Therefore, we designed a paradigm to test whether dissociating the timing of leptin from food consumption is able to disrupt FAA, by administering leptin 3.5-hour in advance of scheduled feeding. To simultaneously record intracellular calcium levels (as a proxy of neural activity) in the DMH^LepR^ neurons, we used an adeno-associated virus (AAV) to Cre-dependently express the calcium indicator GCaMP7s in DMH of LepR-Cre mice (Fig 3A-B, Supplemental Fig 3A-B) ^62^. When these animals were put on SF, we observed robust FAA development by day 3 in the saline control group, which was significantly suppressed by leptin administration (Fig 3C-D). Concurrently, DMH^LepR^ neurons rapidly increased their calcium signal at the time of food delivery, reproducing previous findings and confirming functionality of our system (Supplemental Fig 3C-D) ^52^. To evaluate the data on a circadian timescale, we extracted two readouts from the multi-day fiber photometry calcium recordings (Fig 3B): 1. The “tonic calcium signal” which is the overall intensity of the fluorescence normalized to 24-hour moving average and, 2. The “phasic calcium signal” which is the acute calcium signal increases above the baseline during each recording session as a proxy of dynamic neuron bursts, (Fig 3B, Supplemental Fig 3A)^62–64^. During SF, both types of calcium readouts from the DMH^LepR^ neurons developed responses that predicted the meal time (Fig 3E-O). Specifically, the overall fluorescence (tonic calcium signal) developed an anticipatory decrease prior to food access (Fig 3E-F, I), which is in line with the documented anorexigenic role of DMH^LepR^ neurons ^52,55,56^. Surprisingly, this dampened tonic calcium signal was not impaired by leptin injection but was further potentiated (Fig 3G-I). In contrast, the phasic calcium signal increased during the FAA window in saline injected control animals which was absent in leptin treated mice, in line with the suppressed development of FAA (Fig 3C-D and J-O, Supplemental Fig 3D-F).

**Figure 3.**
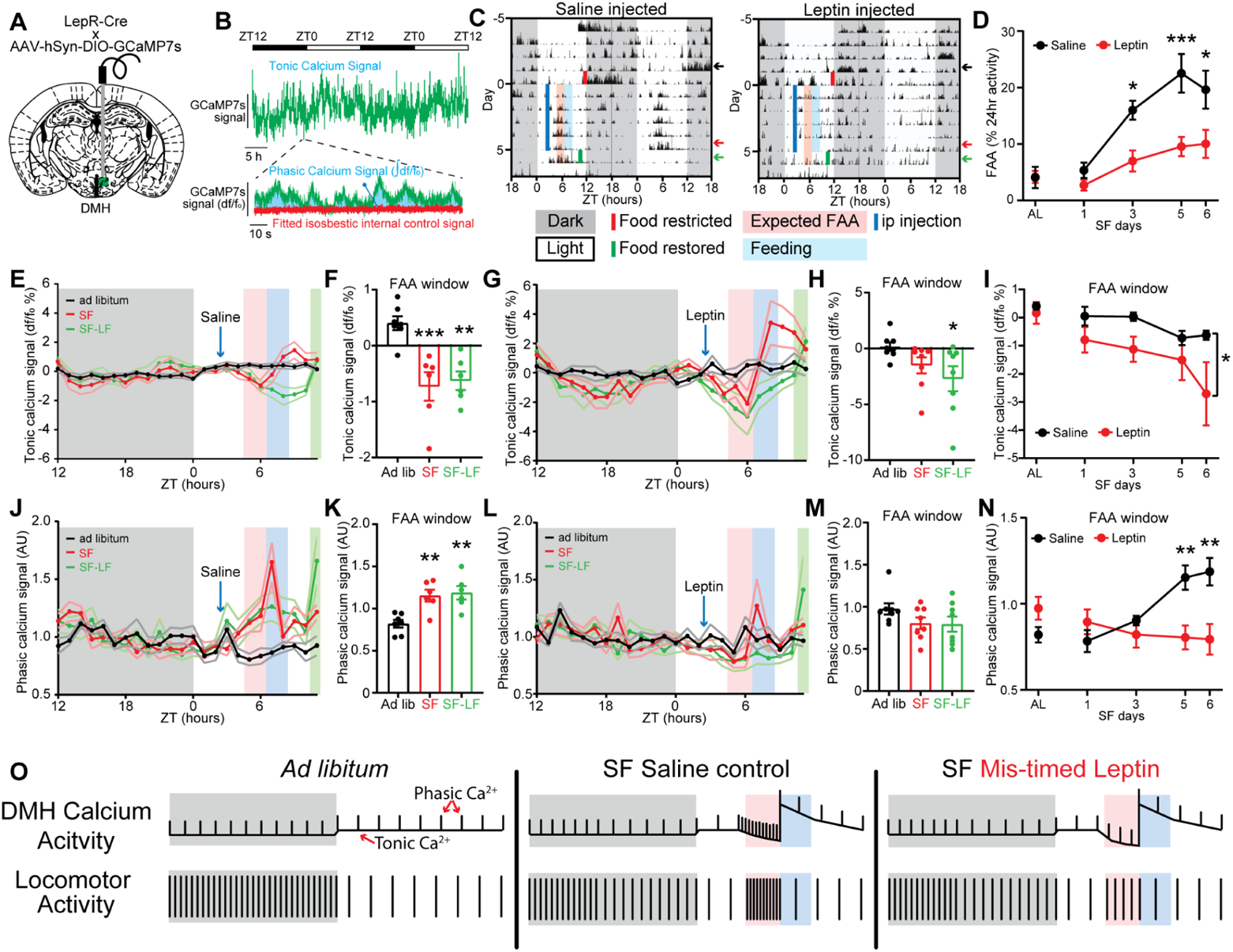
-Leptin sensitive DMH^LepR^ neurons exhibit food entrainable calcium activity patterns which correlate with FAA. **A.** Schematic diagram illustrating unilateral injection of AAV-hSyn-DIO-GCaMP7s and fiber optic cannula implantation to the DMH of LepR Cre mice. **B.** Example data trace illustrating two readouts of long-term fiber photometry calcium imaging. “Tonic calcium signal” represents the total fluorophore brightness. “Phasic calcium signal” represents the intracellular calcium activity over baseline activity in a given recording session. **C.** Representative locomotor actogram of single animals treated with saline (left) or leptin (right) during DMH^LepR^ neuron GCaMP7s fiber photometry recording. Mice are housed in 12:12 LD, fasted at lights off on day 5 (solid red line), injected with saline or leptin at 2.5 hours after lights on (ZT2.5, solid blue line), and fed at ZT6 (2 g on days 1 & 2, 2.5 g on remaining days). Red shaded area is the FAA time window 2 hours pre-meal time (ZT4-6), and blue shaded area is the first 2 hours after food delivery (ZT6-8). Food is restored at ZT10 on day 6 of SF. Color coded arrows indicate three days that are selected for quantification in panels E-H. **D.** Quantification of FAA during long-term DMH^LepR^ neuron GCaMP7s recording. FAA is defined as the locomotor activity in the two-hour window prior to food delivery as a percentage of 24-hour activity. AL indicates *ad libitum* condition two days prior to initiation of drug administration. Mixed-effects (REML) analysis with Bonferroni post hoc comparison; n = 5-8 / group; F_treatment_ (1, 52) = 25.44, p<0.001. **E.** Average tonic calcium signal of DMH^LepR^ neurons from saline control group 2 days before SF (black, *ad libitum*), 5th day during treatment (red, SF), and 6th day where saline injection was withheld and food delivery was delayed for 3.5 hours (green, SF-LF: late feeding). **F.** Quantification of the tonic calcium signal from saline treated mice in the FAA window (average of ZT5-6) from (E). Mixed-effects (REML) analysis with Bonferroni post hoc comparison; n = 6-7 / group; F (2, 16) = 12.27, p=0.0006. **G.** Average tonic calcium signal of DMH^LepR^ neurons from leptin group 2 days before SF (black, *ad libitum*), 5th day during treatment (red, SF), and 6th day where leptin injection was withheld and food delivery was delayed for 3.5 hours (green, SF-LF: late feeding). **H.** Quantification of the tonic calcium signal from leptin treated mice in the FAA window (average of ZT5-6) from (G). Repeated measures one-way ANOVA with Bonferroni post hoc comparison; n = 8 / group; F (2, 14) = 3.596, p=0.0549. **I.** Quantification of the development of the tonic calcium signal during FAA. AL indicates *ad libitum* condition two days prior to initiation of drug administration. Mixed-effects (REML) analysis with Bonferroni post hoc comparison; n = 6-8 / group; F_treatment_ (1, 13) = 4.744, p=0.0484. **J.** Average phasic calcium signal of DMH^LepR^ neurons from saline control group 2 days before SF (black, *ad libitum*), 5th day during treatment (red, SF), and 6th day where saline injection was withheld and food delivery was delayed for 3.5 hours (green, SF-LF: late feeding). **K.** Quantification of the phasic calcium signal from saline treated mice in the FAA window (average of ZT5-6) from (J). Mixed-effects (REML) analysis with Bonferroni post hoc comparison; n = 6-7 / group; F (2, 10) = 15.01, p=0.0010. **L.** Average phasic calcium signal of DMH^LepR^ neurons from leptin group 2 days before SF (black, *ad libitum*), 5th day during treatment (red, SF), and 6th day where leptin injection was withheld and food delivery was delayed for 3.5 hours (green, SF-LF: late feeding). **M.** Quantification of the phasic calcium signal from leptin treated mice in the FAA window (average of ZT5-6) from (L). Repeated measures one-way ANOVA with Bonferroni post hoc comparison; n = 8 / group; F (2, 14) = 2.508, p=0.1172. **N.** Quantification of the development of the phasic calcium signal during FAA. AL indicates ad libitum condition two days prior to initiation of drug administration. Mixed-effects (REML) analysis with Bonferroni post hoc comparison; n = 6-8 / group; F_treatment * time_ (4, 50) = 8.834, p<0.0001. **O.** Summary diagram illustrating the observation of calcium activity pattern in DMH^LepR^ neurons during SF, in animals treated with saline control or mis-timed leptin. Data are represented as mean ± SEM. *p < 0.05; **p < 0.01; ***p < 0.001; ns, not significant. See also supplemental figure 3.

To ensure that the alteration of anticipatory DMH^LepR^ neuron calcium signal by leptin is due to defective food entrainment, rather than acute inhibition of neural activity, we withheld saline/leptin injections on day 6 and delayed food delivery for 3.5 hours (scheduled feeding-late food, SF-LF). We observed that previously saline-treated mice still showed the anticipatory dampening of tonic and elevation of phasic calcium activity during the FAA window which remained until food delivery. Importantly, in the previously leptin-treated group, DMH^LepR^ neurons did not exhibit elevated phasic calcium signal even in the absence of exogenous leptin administration (Fig 3C-O, Supplemental Fig 3D-F).

In summary (Fig 3O), the baseline (tonic) neuronal activity of DMH^LepR^ neurons decreased in anticipation of scheduled food access, and this was further dampened by pre-meal leptin treatment. However, the anticipatory phasic calcium activity, which likely represents the acute increase in neuronal activity ^24,52,55,56^, was largely abolished by mistimed leptin. We therefore posit that the adaptation of DMH^LepR^ neuronal dynamics to scheduled feeding time contributes to the development of behavioral expression of food entrainment.

### Silencing DMH^LepR^ neurons impairs FAA

To determine whether DMH^LepR^ neurons are necessary for food entrainment, we chose to inhibit neuronal transmission by selectively expressing tetanus toxin (TeTx) in these neurons (Fig 4A-B). As observed in DMH lesion studies or previous DMH^LepR^ neuron silencing efforts, the behavioral circadian rhythmicity of animals was largely abolished even during *ad libitum* feeding under 12:12 LD light cycle (Fig 4C-D, supplemental Fig 3G-H) ^23,24,40^, implying a significant role for DMH^LepR^ neurons in expressing circadian behaviors. Following SF, we observed impaired FAA in the TeTx expressing animals compared to their mCherry controls even after 10 days of SF (Fig 4C-E, supplemental Fig 3G-K). This indicates that DMH^LepR^ neuronal output is essential for proper behavioral entrainment to SF.

**Figure 4-.**
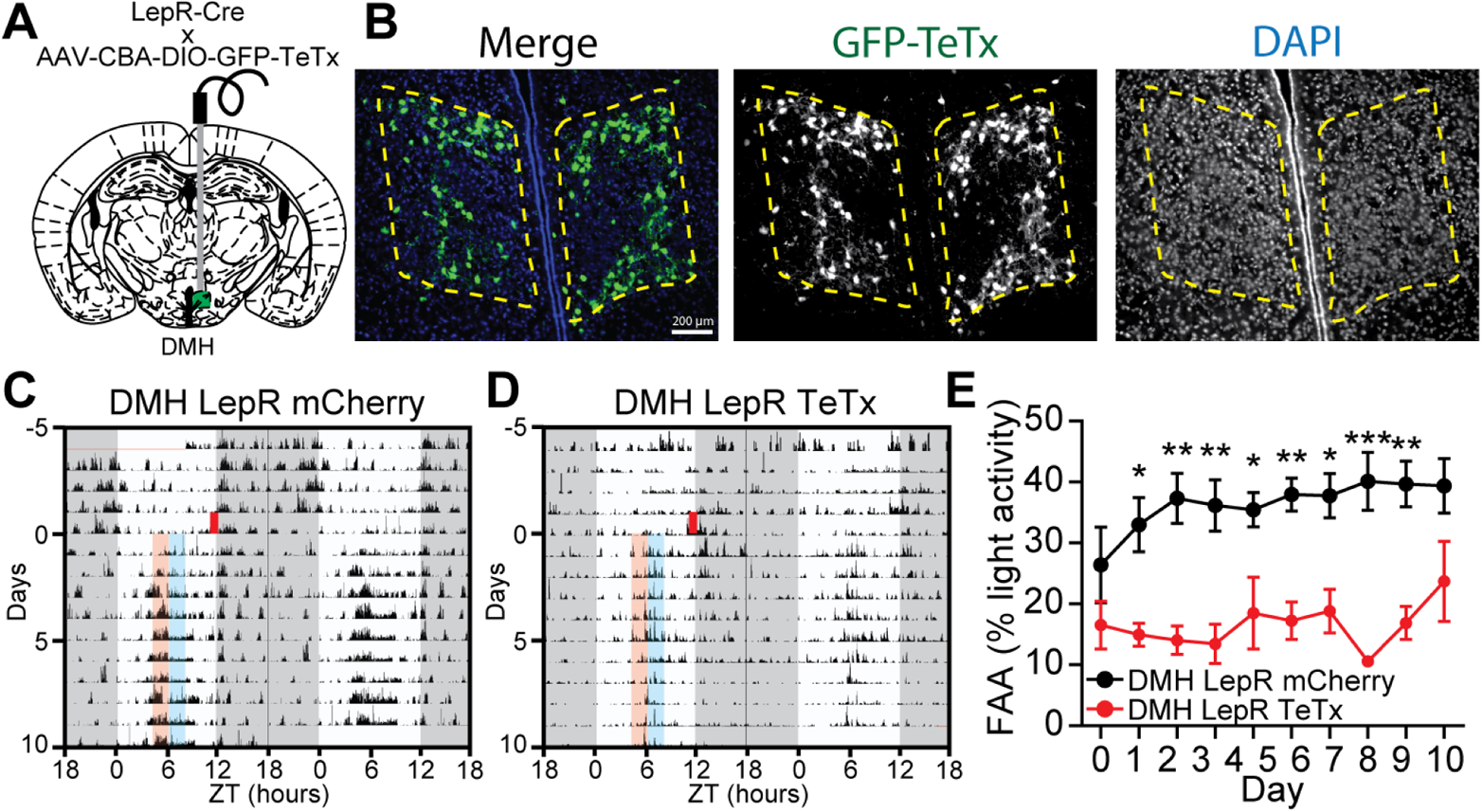
Silencing of DMH^LepR^ neurons impairs FAA. **A.** Schematic diagram illustrating DMH^LepR^ neuron silencing by bilateral injection of AAV-CBA-DIO-GFP-TeTx to the DMH of LepR Cre mice. **B.** Representative images showing the expression of GFP-TeTx in DMH^LepR^ neurons. **C-D.** Representative actograms of (C) DMH^LepR^ mCherry and (D) DMH^LepR^ TeTx mice on SF. Shading color scheme is described in Fig 3C. See supplemental Fig 3 for actograms of all animals. **E.** Quantification of FAA. FAA is defined as the locomotor activity in the two-hour window prior to food delivery as a percentage of light-phase activity. Repeated measures two-way ANOVA with Bonferroni post hoc comparison; n = 5-6 / group; F_virus_ (1, 9) = 33.00, p=0.0003. Data are represented as mean ± SEM. *p < 0.05; **p < 0.01; ***p < 0.001; ns, not significant. See also supplemental figure 3.

### Leptin alters underlying food entrainment pathways, not just appetite

Given the role of leptin as a satiety hormone ^65^, one possibility why leptin blocks the expression of FAA is that its pre-meal treatment suppresses the motivation for the animals to search for food without influencing a timekeeper. Therefore, we designed a cross-over study of wild type mice where we gave saline or leptin for the first 5 days of SF and switched the treatment group on the sixth day for 5 additional days of SF. As in previous experiments, we observed robust FAA by day 3 in the saline control group, while the development of FAA was significantly suppressed by leptin injections (Fig 5A-C, Supplemental Fig 4A-E). On day 6, when the treatment was switched, the animals injected with leptin which previously received saline still exhibited robust FAA compared to those animals injected with saline but previously received leptin (Fig 5A-C). Importantly, this demonstrates that the mistimed leptin is not merely masking entrainment but is instead impairing the establishment of the food timing machinery. Moreover, the FAA of the treatment groups were indistinguishable on Day 7 (Fig 5C) supporting the idea that the food entrainment system is a multi-node network where disrupting one node slows its development but does not eliminate its establishment as the other parts of the system compensate.

**Figure 5-.**
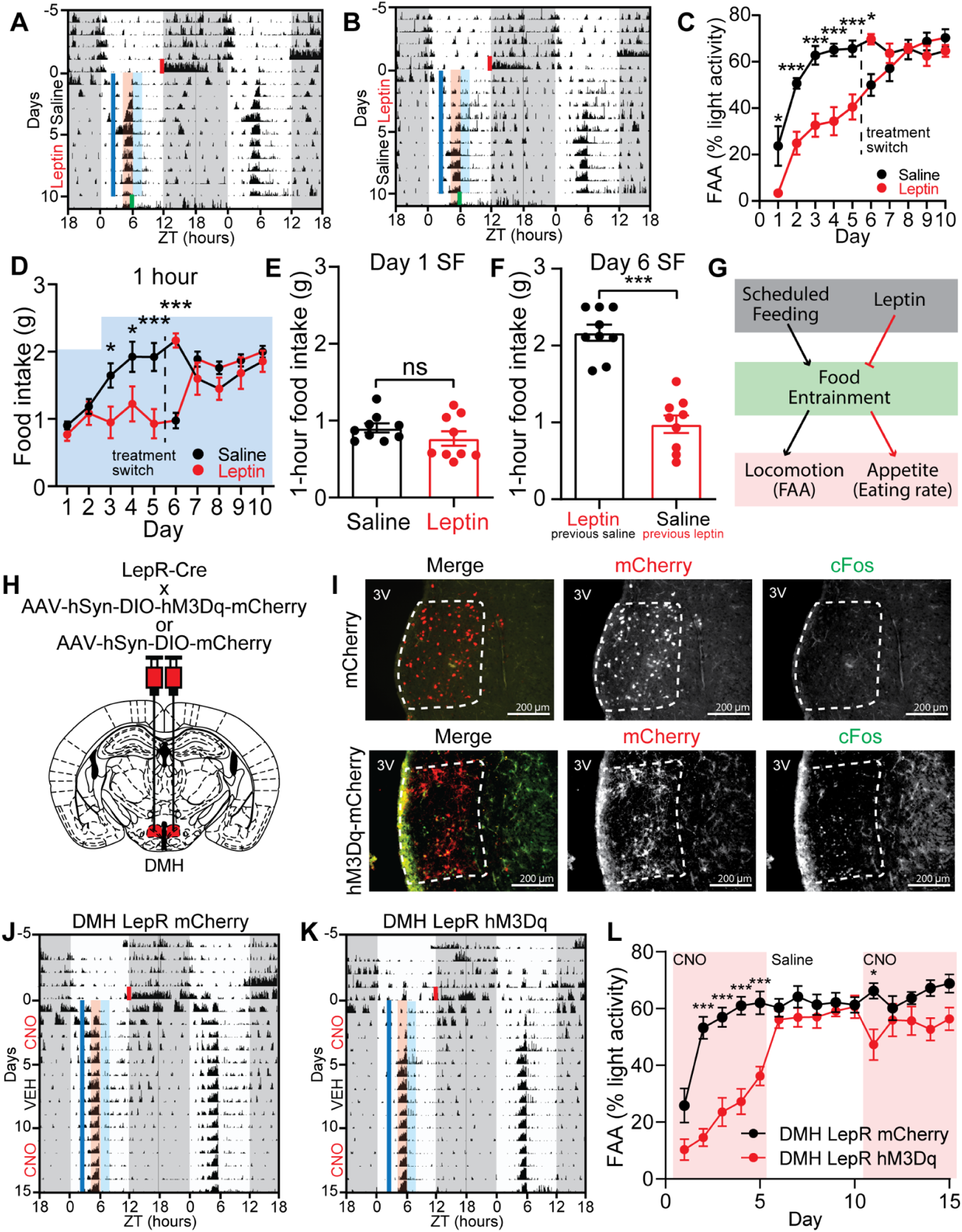
Mis-timed leptin or activation of DMH^LepR^ neurons suppresses the development but not maintenance of food entrainment. **A-B.** Representative actograms of two mice on a 12:12 L:D cycle under *ad libitum* conditions that are then subjected to scheduled feeding (SF) beginning at lights off on SF day 0 and receiving either (A) saline (SF days 1-5) then 5mg/kg leptin (SF days 6-10) or (B) 5mg/kg leptin (SF days 1-5) then saline (SF days 6-10). Shading color scheme is described in Fig 3C. See supplemental Fig 4A-B for actograms of all animals. **C.** Quantification of FAA. FAA is defined as the locomotor activity in the two-hour window prior to food delivery as a percentage of light-phase activity excluding the activity one hour post-injection. Note that red (leptin) or black (saline) data markers indicate the treatment for the day while the data connecting lines identify the initial leptin (red) or saline (black) treatment groups. Repeated measures two-way ANOVA with Bonferroni post hoc comparison; n = 8-9 / group; F_treatment_ (1, 15) = 22.96, p<0.001. **D.** Food intake of mice during SF 1 hour after food delivery. Blue shading indicates the total amount of food that was available for mice to consume on each day. Repeated measures two-way ANOVA with Bonferroni post hoc comparison; n = 9 / group; F_time*treatment_ (9, 144) = 10.15, p<0.0001. **E-F.** First hour food intake on day 1 (E) and day 6 (F) from D. n = 9 / group, Unpaired Student’s t test. Notably, the same treatment groups are illustrated by the same color on the same side of the graphs. **G.** Proposed model illustrating the suppressive effect of leptin on food entrainment, which in turn leads to both decreased locomotion and appetite. **H.** Schematic diagram illustrating bilateral injection of AAV-hSyn-DIO-hM3Dq-mCherry or AAV-hSyn-DIO-mCherry to the DMH of LepR Cre mice. **I.** Representative images showing the expression of mCherry (top) or hM3Dq-mCherry (bottom) in DMH^LepR^ neurons and c-Fos response 2 hours after CNO injection. **J-K.** Representative actograms of (J) DMH^LepR^ mCherry and (K) DMH^LepR^ hM3Dq mice on SF that received 0.3mg/kg CNO (SF days 1-5), saline (SF days 6-10), and 0.3 mg/kg CNO (SF days 11-15) injection at ZT2.5. Shading color scheme is described in Fig 3C. See supplemental Fig 5 for actograms of all animals. **L.** Quantification of FAA. Pink shading indicates days with CNO injection. No shading indicates saline injection. Repeated measures two-way ANOVA with Bonferroni post hoc comparison; n = 8-10 / group; F_virus_ (1, 16) = 15.98, p=0.0010. Data are represented as mean ± SEM. *p < 0.05; **p < 0.01; ***p < 0.001; ns, not significant. See also supplemental figure 4, 5 and 6.

Another key observation we made during the cross-over experiment was that the delayed development of FAA correlated with a slower rate of food consumption, implying that food entrainment prepares the animals for effective foraging during SF (Fig 5D-F). To determine whether the rate of food intake is a distinct output of entrainment and not a prerequisite for the development of FAA, we limited the rate of food consumption during the SF window in a separate cohort of wildtype mice. This paradigm allowed for robust FAA development demonstrating that the rate of food consumption is an additional behavioral output reflecting the strength of food entrainment (Supplemental Fig 4F-I). Results presented here demonstrate that the observed attenuation in food entrainment behaviors in mice injected with leptin during early scheduled feeding (days 1-5) is due to impaired circadian food timing mechanisms rather than acute effects of leptin (Fig 5G).

### DMH^LepR^ neuron stimulation suppresses the development of FAA

Although FAA is largely impaired when the DMH^LepR^ neurons are silenced by the selective expression of TeTx (Fig 4C-E, supplemental Fig 3G-K), these animals are also behaviorally arrhythmic during *ad libitum* conditions in the LD cycle (Fig 4C-D, supplemental Fig 3G-H). To avoid this confound, we took an acute neuronal activity manipulation strategy by using the DREADD hM3Dq to stimulate DMH^LepR^ neurons 3.5 hours before scheduled food time (Fig 5H-I). Similar to leptin treatment, we observed that premature activation of DMH^LepR^ neurons with clozapine N-oxide (CNO) injection impairs the development of FAA over the first 5 days of scheduled feeding. However, it does not significantly impair the maintenance of FAA when CNO is administered after establishment of FAA (Fig 5J-L, Supplemental Fig 5).

These observations demonstrate that acute DMH^LepR^ neuron activation is not masking the behavioral outputs of food entrainment, but instead is attenuating its development. This is in contrast to previous work where chemogenetic inhibition or activation of DMH^Ppp1r1^^7^ neurons during SF showed no effect on FAA ^45^ and in agreement with our snRNAseq findings that DMH^Ppp1r17^ neurons are not responsive to circadian entrainment (Fig 2). Of note, DMH^Pdyn^ neurons have been shown to entrain to SF ^43^ and to reduce the robustness of FAA when silenced ^44^. However, chemogenetic activation of DMH^Pdyn^ neurons only slightly suppressed FAA, which was mainly exhibited after the establishment of food entrainment (Supplemental Fig 6). This underscores that DMH^LepR^ neurons are unique in their role during anticipation of a meal and their precisely timed activity is required for the development of FAA.

### DMH^LepR^ neuron stimulation alters landscape of circadian locomotor activity of *ad libitum* fed mice

We next sought to characterize the impact of DMH^LepR^ neuron acute activation on general circadian behavior in energy replete conditions. To this end, we stimulated the DMH^LepR^ neurons of *ad libitum fed* mice in the middle of the light phase of 12:12 LD cycle, which acutely induced bouts of locomotor activity (Fig 6A-E, Supplemental Fig 7A-C), in line with the previous literature ^66^. Interestingly, we observed a previously undescribed decrease in nighttime locomotor activity compared to controls (Fig 6A-B, D, Supplemental Fig 7A-C). Once injections were stopped after 10 days, daytime activity returned to normal immediately, though it took several days for nighttime activity to fully normalize (Fig 6C-F). To further parse the effects of DMH^LepR^ neuron stimulation on circadian locomotor activity, we repeated this experiment under *ad libitum*, constant dark conditions to remove the entraining effect of light. Strikingly, in constant darkness DMH^LepR^ stimulation partitioned circadian locomotor activity without changing the free-running period (Fig 6G-O, Supplemental Fig 7D-G). Activation early in the active phase (~circadian time [CT]14) advanced the offset of locomotor activity to approximately 3 hours after CNO injection, with a concomitant reduction in 24-hour activity (Fig 6G-J, Supplemental Fig 7D-E). Activation late in the active phase (~CT22) delayed the offset of locomotor activity to approximately 2 hours following CNO injection, with no change in total locomotor activity compared to controls (Fig 6K-O, Supplemental Fig 7F-G). To our surprise, this delayed running wheel activity in DMH^LepR^ hM3Dq mice persisted after cessation of CNO injections, in phase with the previous day’s injection time (Fig 6K-O). These data demonstrate that activation of DMH^LepR^ neurons can partition and entrain circadian locomotor activity in a time-dependent manner, and suggests DMH^LepR^ neurons are an integrating hub that connects the food entrainable clock with the light entrainable clock.

**Figure 6-.**
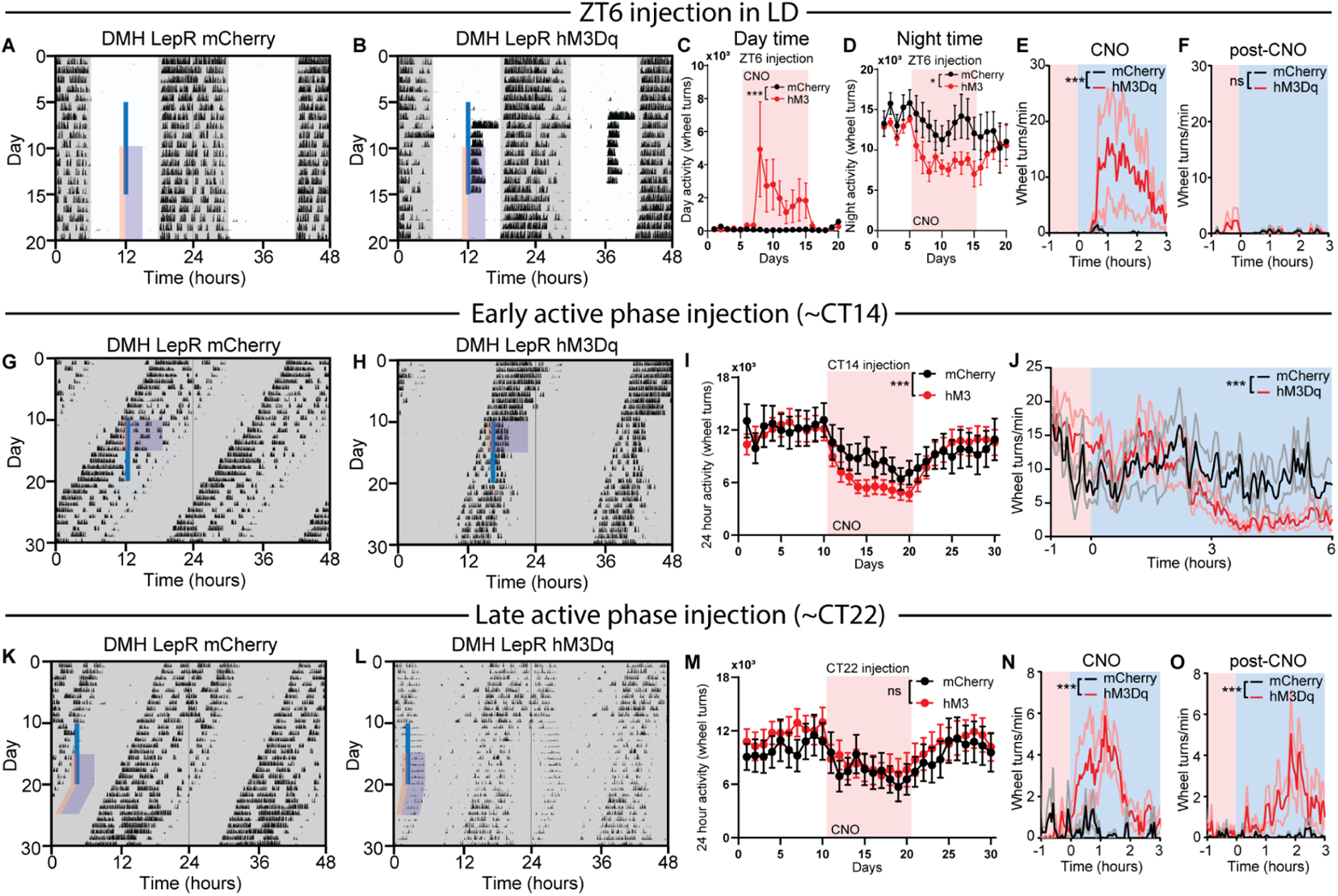
Repetitive activation of DMH^LepR^ neurons alters circadian locomotor activity. **A-B.** Representative actograms of LepR Cre animals bilaterally injected with (A) AAV-hSyn-DIO-mCherry or (B) AAV-hSyn-DIO-hM3Dq-mCherry and injected with 0.3mg/kg CNO at ZT6 (solid blue line) in 12-12 LD with access to a running wheel. Pink shading represents one hour prior to injection, and blue shading 3 hours after injection. See supplemental Fig 7A-B for actograms of all animals. **C.** Light phase (Day) wheel revolutions of DMH^LepR^ mCherry and DMH^LepR^ hM3Dq animals before, during, and after CNO injections at ZT6 in 12-12 LD. Pink shading represents days with CNO injection. Repeated measures two-way ANOVA; n = 5-6 / group; F_virust * time_(19, 171) = 3.302, p<0.001. **D.** Dark phase (Night) wheel revolutions of DMH^LepR^ mCherry and DMH^LepR^ hM3Dq animals before, during, and after CNO injections at ZT6 in 12-12 LD. Pink shading represents days with CNO injection. Repeated measures two-way ANOVA; n = 5-6 / group; F_virust * time_(19, 171) = 1.707, p=0.0390. **E.** Average wheel running activity induced by chemogenetic activation of DMH^LepR^ neurons at ZT6 in 12-12 LD. Color coded time window is indicated in (A-B). Repeated measures two-way ANOVA; n = 5-6 / group; F_virust * time_(79, 711) = 2.519, p<0.001. **F.** Quantification of activity for 5 days after cessation of CNO injections. Color coded time window is indicated in (A-B). Repeated measures two-way ANOVA; n = 5-6 / group; F_virus_ (1, 9) = 0.9519, p=0.3547; F_virus*time_ (79, 711) = 1.202, p=0.1215. **G-H.** Representative actograms of LepR Cre animals bilaterally injected with (G) AAV-hSyn-DIO-mCherry or (H) AAV-hSyn-DIO-hM3Dq-mCherry and injected with 0.3mg/kg CNO at ~CT14 (solid blue line), in *ad libitum*, constant dark conditions with access to a running wheel. Pink shading represents one hour prior to injection, and blue shading 6 hours after injection. See supplemental Fig 7D-E for actograms of all animals. **I.** 24-hour total wheel revolutions of DMH^LepR^ mCherry and DMH^LepR^ hM3Dq animals before, during, and after CNO injections at ~CT14. Animals were housed under *ad libitum*, constant dark conditions. Pink shading represents days with CNO injection. Repeated measures two-way ANOVA; n = 9-12 / group; F_virust * time_ (29, 551) = 2.468, p<0.001. **J.** Average wheel running activity induced by chemogenetic activation of DMH^LepR^ neurons at ~CT14. Color coded time window is indicated in (E-F). Repeated measures two-way ANOVA; n = 9-12 / group; F_virust * time_(140, 2660) = 2.114, p<0.001. **K-L**. Representative actograms of LepR Cre animals bilaterally injected with (K) AAV-hSyn-DIO-mCherry or (L) AAV-hSyn-DIO-hM3Dq-mCherry and injected with 0.3mg/kg CNO at ~CT22 (solid blue line), in *ad libitum*, constant dark conditions with access to a running wheel. Pink shading represents one hour prior to injection, and blue shading 3 hours after injection. See supplemental Fig 7F-G for actograms of all animals. **M.** 24-hour total wheel revolutions of DMH^LepR^ mCherry and DMH^LepR^ hM3Dq animals before, during, and after CNO injections at ~CT22. Animals were housed under *ad libitum*, constant dark conditions. Pink shading represents days with CNO injection. Repeated measures two-way ANOVA; n = 9-12 / group; F_virus_ (1, 19) = 0.2061, p=0.6550; F_virus*time_ (29, 551) = 0.7120, p=0.8676. **N.** Average wheel running activity induced by chemogenetic activation of DMH^LepR^ neurons at ~CT22. Color coded time window is indicated in (K-L). Repeated measures two-way ANOVA; n = 9-12 / group; F_virus_ (1, 18) = 8.580, p=0.0090; F_virus*time_ (80, 1440) = 2.029, p<0.001. **O.** Quantification of sustained activity for 5 days after cessation of CNO injections. Color coded time window is indicated in (K-L). Repeated measures two-way ANOVA; n = 8-12 / group; F_virus*time_ (80, 1440) = 1.745, p<0.001. Data are represented as mean ± SEM. *p < 0.05; **p < 0.01; ***p < 0.001; ns, not significant.

### DMH^LepR^ neurons link metabolic and circadian systems via direct projection to the SCN

Finally, we sought to determine if the DMH^LepR^ neurons are directly communicating metabolic signals to the primary circadian pacemaker in the SCN. We first asked whether the SCN was necessary to mediate the effects of DMH^LepR^ neuron activation on circadian locomotor activity. We electrolytically lesioned both sides of the SCN (SCNxx) in mice expressing hM3Dq or mCherry in the DMH^LepR^ neurons (Fig 7A-C), and performed the 10-day chemogenetic activation protocol in constant darkness with *ad libitum* food. We observe an acute bout of wheel running activity in SCNxx DMH hM3Dq, but not SCNxx mCherry control mice after CNO injection (Fig 7D-F, Supplemental Fig 7H-I). In contrast to the SCN-intact experiments, we did not observe persistence of this activity in SCNxx animals following cessation of CNO injection on subsequent days (Fig 7G, Supplemental Fig 7H-I). This finding suggests that DMH induced locomotor activity is independent of the SCN, however, the SCN is necessary for maintaining DMH^LepR^ induced circadian behavior.

**Figure 7-.**
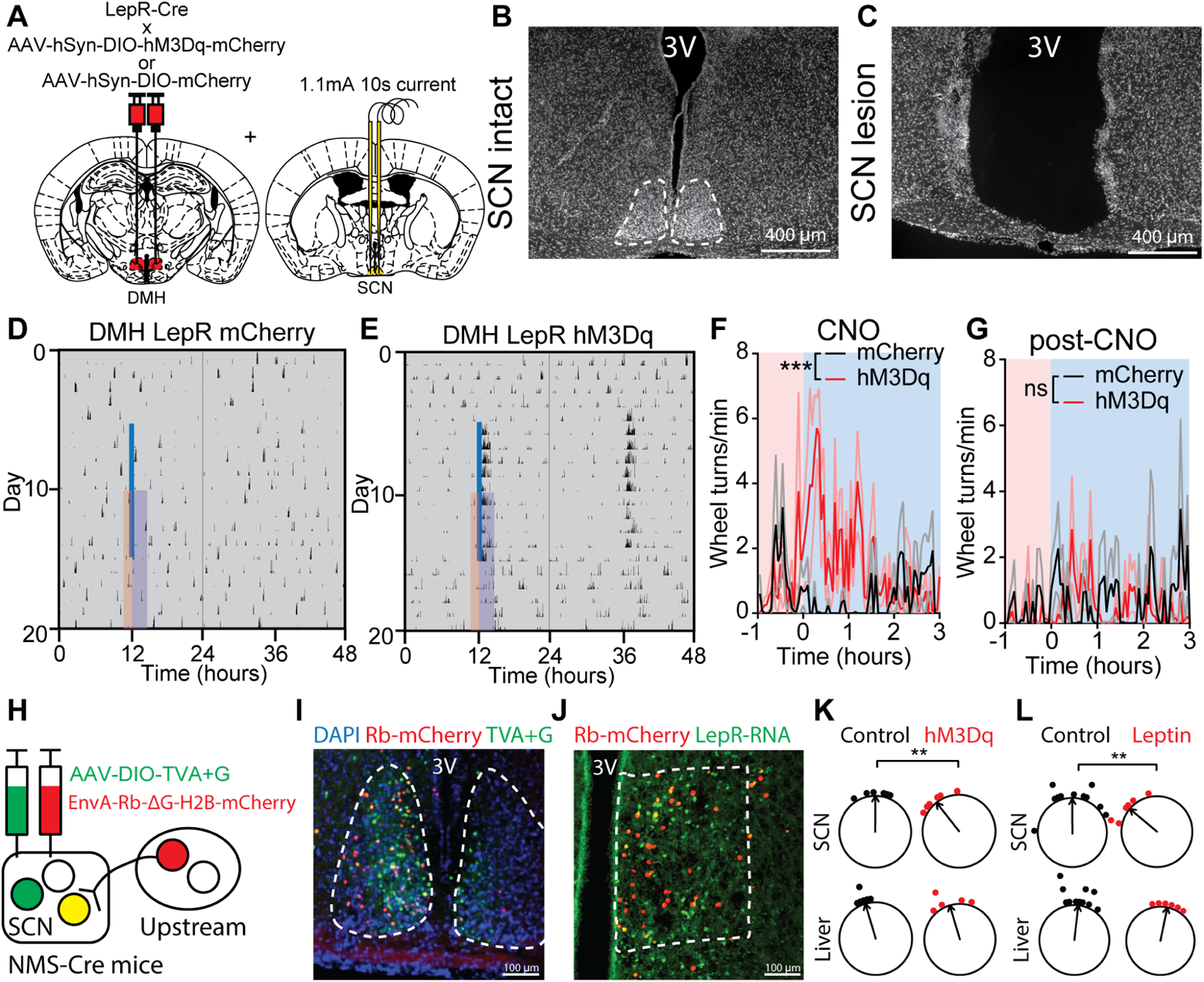
DMHLepR neurons alter circadian behavior via the SCN. **A.** Schematic diagram illustrating bilateral injection of AAV-hSyn-DIO-hM3Dq-mCherry or AAV-hSyn-DIO-mCherry to the DMH of LepR Cre mice coupled with electrolytic lesioning of the SCN in the same mice. **B-C.** Representative DAPI staining images of (B) intact and (C) electrolytic lesioned SCN. **D-E.** Representative actograms of an SCN lesioned DMH^LepR^ (D) mCherry, and (E) DMH^LepR^ hM3Dq animal injected with CNO every 24 hours for 10 days in constant darkness. Solid blue line indicates CNO injection. See supplemental Fig 7H-I for actograms of all animals. **F.** Average wheel running activity induced by chemogenetic activation of DMH^LepR^ neurons from (D-E). Color coded time window is indicated in (D-E). Repeated measures two-way ANOVA; n = 5-6 / group; F_virus_ (1, 9) = 8.245, p=0.0184; F_virust*time_ (79, 711) = 2.432, p<0.0001. **G.** Quantification of activity for 5 days after cessation of CNO injections. Color coded time window is indicated in (D-E). Repeated measures two-way ANOVA; n = 5-6 / group; F_virus_ (1, 9) = 0.3450, p=0.5714; F_virus*time_ (79, 711) = 1.047, p=0.3746. **H.** Schematic illustration of NMS Cre dependent rabies virus monosynaptic retrograde tracing. **I-J.** Representative images of (I) SCN and (J) DMH from retrograde tracing strategy in H. **K.** The ZT phase of the first bioluminescence peak of SCN and Liver from PER2::luciferase mice injected with CNO at ZT6 (Control), or PER2::luciferase;LepR-Cre mice with hM3Dq expressed in DMH^LepR^ neurons and injected with CNO at ZT6 (hM3Dq). Two-way ANOVA with Bonferroni post hoc comparison; n = 5-6 / group; F_treatment_(1, 19) = 5.361, p=0.0319. **L.** The ZT phase of the first bioluminescence peak of SCN and Liver from control (mixture of ZT6 saline or untreated) or ZT6 leptin injected PER2::luciferase mice. Two-way ANOVA with Bonferroni post hoc comparison; n = 6-11 / group; F_treatment_ (1, 30) = 4.772, p=0.0369. Control group is the same dataset as in fig 1B, re-plotted and analyzed with ZT6 leptin treated animals. Data are represented as mean ± SEM. *p < 0.05; **p < 0.01; ***p < 0.001; ns, not significant.

While the DMH makes reciprocal connections with SCN neurons via direct and indirect projections, it is unknown whether DMH^LepR^ neurons are directly connected ^67^. To identify potential direct DMH^LepR^ projections, we carried out Cre-dependent monosynaptic retrograde viral rabies tracing, and saw that DMH^LepR^ neurons innervate SCN neuromedin S (NMS) neurons (Fig 7H-J). To functionally test this connectivity, we selectively activated DMH^LepR^ neurons at ZT6 for 4 days using hM3Dq in PER2LUC mice, and observed a phase advance in the SCN PER2 rhythm similar to animals on SF at ZT6. However, this stimulation did not induce a phase shift in the liver (Fig 7K). Similarly, we observed that 4 days of leptin injection at ZT6 significantly phase advanced the SCN but not the liver (Fig 7L). These data point to the existence of a leptin→DMH^LepR^→SCN axis which communicates the metabolic status to the central circadian system.

## Discussion

The circadian system synchronizes to salient non-photic cues, such as timed availability of food, receptive mates, or exercise ^11,68,69^. In this work, we demonstrated that leptin, in combination with one of its central nervous system targets, the DMH^LepR^ neurons, form an essential node that links food intake with the development of circadian food entrainment. In the process, we also identified an intriguing property of the circadian system whereby locomotor activity is partitioned into at least two components in response to overactivation of the DMH^LepR^ neurons. By functional direct innervation to the SCN, DMH^LepR^ neurons, have the potential to serve as an information conversion point for non-photic entrainment. The methodological paradigms presented here offer a new platform to test the involvement of other molecular signals and anatomic regions in the development and maintenance of food entrainment, and the relative function of inputs to the circadian system.

### Impact of SF on canonical clock genes in the DMH

In our snRNAseq analysis of the DMH, we did not observe a significant difference in the KEGG annotated “circadian rhythm” pathway (canonical circadian genes, e.g. *Bmal1*, *Clock*; https://www.kegg.jp/entry/map04710) except for *Per3* expression (Supplemental Fig 2). However we cannot rule out an alteration in the expression of other core circadian genes, since they possess different phases, which might require analysis at multiple time points across the day to reveal a change. In contrast, we did identify genes in the “circadian entrainment” pathway to be altered in the DMH (Fig 2, Supplemental Fig 1I, 2D). Although this pathway was originally annotated based on the studies of light entrainment of SCN, DMH neurons likely share similar machinery to alter their own molecular rhythms.

### Parameters of food entrainment

In addition to FAA, food entrainment is exhibited by other behaviors and physiological processes, sometimes with differential outcomes (Fig 1A). For instance, peripheral organs, especially the liver, are more susceptible to food entrainment with the ketone body β-hydroxybutyrate as a key regulator ^21,30,70^. While FAA is a widely used central nervous system driven behavioral output of food entrainment, we characterized the rate of food consumption as an additional readout that has not received as much attention. Furthermore, we quantified food entrainment at two stages: development and maintenance. While the speed of entrainment to light is a widely accepted way to study the strength of light entrained clock ^71–73^, except for a few publications ^49,72,74^, food entrainment is exclusively gauged by existence of FAA, or robustness of FAA at the maintenance stage. Our work highlights the importance of development stage of food entrainment and places leptin and DMH^LepR^ neurons as a part of the food clock network.

### Leptin, DMH^LepR^ neurons, and their connections coordinate food entrainment

Previous findings regarding the role of leptin circadian regulation have been mixed. While leptin has been shown to advance the SCN when administered directly to brain slices of rats ^75^ or *in vivo* in mice (Fig 7L), it does not appear to appreciably alter circadian behavior in mice ^76^. However, leptin suppresses FAA when administered continuously to leptin-deficient mice ^77,78^, or rats in an activity-based anorexia model ^79^. The data presented here lend further credence to the importance of the timing of leptin and position it as a putative circadian entraining agent. This highlights the need for future work investigating the role of mistimed leptin release as a result of snacking outside of mealtimes contributing to the adverse metabolic outcomes of modern eating habits ^80^.

Leptin appears to exert at least some of its effects via the DMH, a hypothalamic region that serves as an anatomic integration hub to control food intake, thermogenesis, locomotor activity, and circadian behaviors ^24,52,66,81,82^. Interestingly, in addition to the DMH^LepR^-SCN circuit described in this work, at least one other target of the DMH^LepR^ neurons, the AgRP neurons in the ARC, have been implicated in the development of FAA ^83,84^. Therefore, we speculate that the DMH, via at least its projections to SCN and AgRP neurons, serves as a critical node for food entrainment by coordinating circadian and appetitive behaviors. This connectivity accommodates the previously demonstrated redundancy in the food entrainable network, as it has been convincingly shown that none of these individual regions are necessary for eventual development of food entrainment ^12,13,83, 85–87^. However, ablation of one or more components does change the quality of food entrainment behaviors.

### Role of DMH^LepR^ neurons within the central pacemaker

We demonstrated that the DMH^LepR^ neurons cannot solely be an output for FAA as its manipulation does not inhibit food entrainment once established (Fig 7). Additionally, scheduled activation of DMH^LepR^ neurons in constant conditions induces a partitioned secondary bout of activity that remains phase-locked to the primary onset of activity even after cessation of the stimulation (Fig 6). These observations led us to postulate the existence of at least two coupled but independent clocks in the DMH and SCN akin to the “morning” and “evening” oscillators proposed in *Drosophila* ^88^. However, it remains to be determined if the DMH and SCN are the two coupled clocks that can function independently in certain conditions, or if DMH partially uncouples the SCN which harbors these two clocks.

It has been increasingly acknowledged that when we eat, in addition to what and how much, plays a critical role in maintaining metabolic homeostasis and health ^4–6,8,89–91^. This bidirectional relationship between food and the circadian clock leads to dysfunction of both the circadian system and metabolic homeostasis. Further understanding of the circuits that govern these food-circadian interactions will provide new avenues to improve metabolic health ^9,92–94^.

## AUTHOR CONTRIBUTIONS

Q.T., E.G., C.D.D, B.P., J.N.C., and A.D.G. conceived and designed the experiments, and wrote the manuscript with input from all co-authors. Q.T., E.G., and B.P. performed the generation of experimental cohorts, data collection, analysis and interpretation. Specifically, Q.T. performed calcium imaging. E.G. performed RNA FISH staining. B.P. made the initial observations of leptin and DMH^LepR^ activation’s behavioral effects. E.G. and R-J.A-F. conducted the snRNAseq experiments, whereas E.G. analyzed snRNAseq data. C.D.B. and Q.Z. contributed to intracranial surgery. R.O. and T.B.G. contributed to body weight and food intake measurements during SF. Q.Z. and S.P.W. contributed to DMH^LepR^ neuron chemogenetic activation in constant darkness. C.D.B., Q.Z., S.P.W, T.B.G., R.O., R.S., and J.O. contributed to daily animal husbandry during long-term circadian behavior experiments.

## ACKNOWLEDGEMENTS

We thank the members of the Güler, Deppmann, Campbell, and Provencio labs (University of Virginia), Yamazaki lab (University of Texas Southwestern Medical Center) and Steele lab (California State Polytechnic University, Pomona) for comments and suggestions on the preparation of the manuscript. We are thankful for technical advice from Austin Byler Keeler, Jianhua Cang and Chad Daniel Meliza (electrolytic lesion, University of Virginia), and Pantong Yao (*in vivo* calcium imaging data interpretation, University of California San Diego). Additionally, we are grateful for the access to the confocal microscope generously provided by Cang and Liu labs (University of Virginia). We are grateful for the generous gift of the cre dependent TeTx virus from Zweifel lab (Washington University). We thank the University of Virginia School of Medicine Genome Analysis and Technology Core, RRID:SCR_018883 for sequencing our samples and quality control. This work was supported by NIH T32-GM7267-39 and NIH T32-GM7055-45 (B.P.), NIH R01GM121937 and R35GM140854 (A.D.G.), NIH R01HL153916-01A1 and American Diabetes Association Pathway to Stop Diabetes award 1-18-INI-14 (J.N.C.), and a UVA Wagner Fellowship (B.P.).

## RESOURCE AVAILABILITY

Raw snRNAseq files can be found at gene expression omnibus (GEO) with accession code GSE211757.

## METHODS

### Single Nucleus RNA Sequencing (Figs 1-3, Extended data Figs 1-2)

#### Mouse Lines

All experiments were carried out in compliance with the Association for Assessment of Laboratory Animal Care policies and approved by the University of Virginia Animal Care and Use Committee. Animals were housed on a 12-h light/dark cycle with food (PicoLab Rodent Diet 5053) and water *ad libitum* unless otherwise indicated. For generation of the 10X single nucleus RNA-seq data (Figs 1-3, Extended data Figs 1-2), we used both male and female LepR-cre mice (B6.129-Lepr^tm3(cre)Mgmj^/J, The Jackson Laboratory #032457, RRID:IMSR_JAX:032457) ^95^ crossed to Ai14 tdTomato reporter line (B6.Cg-*Gt(ROSA)26Sor^tm^*^14*(CAG-tdTomato)Hze*^/J, Strain #007914, RRID:IMSR_JAX:007914) ^96^.

#### Scheduled Feeding in Comprehensive Lab Animal Monitoring System (CLAMS)

Indirect calorimetry in the CLAMS system (Columbus Instruments) was used to evaluate metabolic parameters and ambulatory locomotor activity during *ad libitum,* overnight fasted, or time and calorie restricted scheduled feeding (SF). All mice were on a 12:12 light-dark cycle with *ad libitum* access to food and water unless otherwise indicated. Lepr-cre; tdTomato mice were singly housed and acclimated to the CLAMS for 3 days prior to experiment start. The night before SF began, the SF cohort was fasted and cages were changed and thereafter were given 2-3 grams of food (PicoLab Rodent Diet 5053) at zeitgeber time (ZT) 6 for 10 days. The overnight fasted cohort had *ad libitum* food access until the night before sacrifice, when food was removed and cages were changed. We repeated this experiment a total of 3 times with 2 mice per condition each time (n = 6 mice / condition total).

#### Brain Extraction and Microdissection

All mice were sacrificed at ZT 5. Brains were immediately extracted and dropped into ice cold Hanks’ Balanced Salt Solution (HBSS). After 2 minutes, brains were embedded in low melting point agarose (Precisionary Instruments, Natick, MA) and sectioned at 400 µm on a Compresstome VF-200 Vibrating Microtome (Precisionary Instruments, Natick, MA, USA) into DNAse/RNAse free 1x PBS. Hypothalamic sections of interest were immediately collected into RNAprotect (Qiagen, Hilden, Germany) and kept in RNAprotect at 4°C overnight. The next day Lepr-Cre; TdTomato positive cells were visualized using a fluorescent stereoscope (Leica, Wetzlar, Germany). tdTomato fluorescence was used to approximate DMH and SCN boundaries during microdissection of these regions. DMH and SCN microdissected tissue samples were placed into Eppendorf tubes separated by brain region and feeding condition, and stored in −80°C until nuclei isolation.

#### Isolation of Single Nucleus for RNA sequencing

DMH and SCN microdissected tissue samples were transferred from a −80°C freezer and into individual 2mL glass dounce homogenizer tubes (Kimble, Vineland, NJ, USA) to be homogenized according to a protocol modified from the method described previously ^97^. Tissue was homogenized in 1mL of buffer (16mM sucrose, 5mM CaCl, 3mM Mg(Ac)_2_, 10mM Tris pH 7.8, 0.1mM EDTA, 1% NP40, 1mM beta-mercaptoethanol in H_2_O) on ice, with 25 passes of pestle A and 25 passes of pestle B. An additional 4mL of buffer was added to the nuclei suspension and placed on ice for 5 minutes. Then 5 mL of 50% OptiPrep Density Gradient Medium [Sigma Aldrich, MO, USA] (30mM CaCl, 18mM Mg(Ac)_2_, 60mM Tris pH 7.8, 0.1 mM PMSF, 6mM beta-mercaptoethanol in H_2_O; in 60% (w/v) solution of iodixanol in sterile water) was added to nuclei on ice and inverted 10 times to mix. The nuclei suspension was layered onto 10 mL of 29% OptiPrep solution (Buffer + 50% OptiPrep) in a 38.5 mL Ultra-Clear tube (Beckman-Coulter, CA, USA) before being centrifuged at 7,333g for 30 minutes at 4°C. Supernatant was discarded and the nuclei pellet was resuspended in 1x PBS + 1% BSA (Sigma-Aldrich, MO, USA) + 2 mM Mg^2+^ + 0.1% RNase inhibitor (Sigma-Aldrich, MO, USA) for 15 minutes on ice. The nuclei suspension was pipetted through a 20 µm mesh filter along with 2 drops of propidium iodide (PI) Ready Flow reagent (Thermo Fisher, MA, USA) and immediately taken on ice to be FACS sorted using an SH800 (Sony, Tokyo, Japan) cell sorter. Sorting was gated to select for PI+ single nucleus. Nuclei were sorted through a 70 µm nozzle into a 2mL LoBind collection tube (VWR, PA, USA) containing 18.8uL of RT Reagent B from the Chromium Next GEM Single Cell 3’ Reagent Kit v3.1. The remaining components of the 10X Step 1 mastermix were then gently mixed with the contents of the FACS collection tube and loaded into the 10X Genomics Chromium Controller Chip G.

#### Single-Nucleus RNA-Seq Workflow

The single-nucleus samples were processed into sequencing libraries using the Chromium Next GEM Single Cell 3’ Reagent Kit v3.1 according to the manufacturer’s protocol (version 3.1, revision D). After generation of the GEMs (Gel Bead-In EMulsions) and reverse transcription of poly-adenylated mRNA, cDNA was amplified (10-14 cycles), enzymatically fragmented, and ligated to Illumina adapters. Sequencing libraries were indexed, size selected for 400-600 bp using SPRIselect (Beckman Coulter, Indianapolis, IN, USA), and quantified by Qubit (v4.0, 1X high-sensitivity dsDNA kit, Thermo Scientific) and Bioanalyzer (Agilent, Santa Clara, CA, USA). Size-corrected library concentrations were used to pool libraries for equimolar representation. The pooled concentration was measured by KAPA Library Quant qPCR according to the manufacturer’s instructions (KAPA Biosciences, Wilmington, MA) and by Qubit. Library pools were sequenced using a P100 cycle kit on the NextSeq 2000 (Illumina, CA, USA) in the University of Virginia School of Medicine Genome Analysis and Technology Core, RRID:SCR_018883. The sequencing structure was as follows: Read 1 was 28 bp (16 bp barcode, 12 bp UMI), Read 2 was 98 bp (cDNA) and Index 1 was 8 bp (single index). Overall, we had 7 DMH and 5 SCN sequencing library pools. 10X Batch 1 & 2 contained mixed pools of all feeding conditions, but 10X Batch 3 contained a unique feeding condition per library pool (Extended data Fig. 1A, E).

#### Single-Nucleus RNA-Seq Data Processing

Raw digital expression matrix files for each sequencing run were transferred to a high-performance cluster (HPC) server where they were demultiplexed based on sample index. We generated fastq files with bcl2fastq2 version 2.20.0, then used Cell Ranger version 5.0.0 to align transcripts to the Cell Ranger supplied mouse genome, mm10 2020-A (GENCODE vM23/Ensembl 98), quantify expression levels, and partition them according to their cell-specific barcode. We ran the Cell Ranger count program with the ”--include-introns” argument to include intronic reads in the gene expression quantitation.

#### Single-Nucleus RNA-Seq Analysis

Cell Ranger h5 files were read into Seurat v4 ^98^ in R (version 4.1.0) and RStudio (version 1.4.1717) and merged by brain region (DMH, SCN) for clustering analysis. We filtered the initial datasets to remove low quality samples (i.e., cells with less than 100 genes detected or greater than 0.5% mitochondrial reads). We then log-normalized the data; selected 2,000 most variable genes (“feature selection”), and scaled gene expression. We performed Principal Component Analysis (PCA) to linearly reduce the dimensionality of the highly variable gene set. We defined distance metrics based on K-nearest neighbor analysis, grouped cells with Louvian algorithm modality optimization, and visualized cell embeddings in low-dimensional space with Uniform Manifold Approximation and Projection (UMAP) nonlinear dimensionality reduction. To focus our analysis on neurons, we subsetted neuronal clusters based on their enriched expression of neuronal marker genes (*Syt1*, *Syn1*, *Tubb3*). To correct for batch effects, we integrated across sample batches using Seurat’s function for reciprocal principal component analysis (RPCA). Next, we subsetted region-specific clusters based on expression of positive and negative marker genes for each target brain region (SCN, DMH), as described in the next section. We reclustered the identified DMH and SCN neurons using the following parameters: DMH, 2,000 most variable genes, first 15 PCs, resolution setting of 0.8; SCN, 2,000 most variable genes, first 13 PCs, resolution setting of 0.5 Finally, we assessed cluster markers with the Wilcoxon Rank Sum test using Seurat default settings. Cluster markers were selected based on top p-values (adjusted to correct for multiple comparisons), high percent expression within the cluster and low percent expression outside of the cluster, and validated based on Allen Brain Atlas mouse *in situ* hybridization data and previous literature.

#### Identification of DMH and SCN neurons

DMH neuron types were selected based on previously reported markers ^51^, as well as known highly expressed genes in the DMH including *Lepr, Pdyn, Ppp1r17, Cck* and *Grp* ^45,57,95^. Markers were validated via the Allen Brain Atlas mouse *in situ* hybridization data. Additionally, clusters enriched with genes expressed in surrounding hypothalamic regions but not DMH were excluded from further analysis, including the following: PVH (*Sim1*) ^99^, VMH (*Slit3, Qrfpr, Arpp21, Nr5a1, Fezf1*) ^100^, Arc (*Prlr, Nr5a2*) ^101^, LH (*Pvalb, Klk6, Nts*) ^102,103^, or the tuberomammillary nucleus (*Hdc*) ^104,105^. Cluster markers were prioritized based on multiple-comparison adjusted p-values, high percent expression within the cluster and low percent expression outside of the cluster. We defined SCN neuron populations based on previous literature by plotting expression of cluster specific markers and circadian genes from published datasets by our clusters ^35,36,67^.

#### Functional Analysis of Differentially Expression Genes

To find differential gene expression between feeding condition groups (*ad libitum*, fasted, SF), we used Seurat’s ‘FindMarkers’ function to run Wilcoxon Rank Sum statistical tests. False Discovery Rate (FDR) was calculated using the ‘p.adjust’ function. We set cutoffs of log2FC > 0.25 and FDR < 0.05 to quantify the total number of differentially expressed genes per feeding condition comparison. To visualize STRING functional protein interaction networks, we used Cytoscape open-source software (version 3.9.1). We input lists of differentially expressed genes in the cluster 13_Lepr between both SF and fasted and SF and *ad libitum* using the same criteria as previously described above. We ran KEGG Gene Ontology functional enrichment on the mouse genome to label genes involved in upregulated pathways during SF. We measured differentially expressed pathways among the three feeding conditions using the ‘enrichR’ package. To visualize genes and pathways differing significantly between feeding conditions, we used the ‘DEenrichR’ function which applies the Wilcoxon Rank Sum test to identify differentially expressed (DE) genes (log2(fold change) > 0.25, multiple-comparison adjusted p < 0.05). Significantly DE genes are then scored based on odds ratios to fall into pathways categories defined by the Kyoto Encyclopedia of Genes and Genomes (KEGG) 2019 Mouse database. With the ‘ggplot2’ package, we then graphed upregulated and downregulated pathways in each comparison on a superimposed bar graph, colored by condition specific comparison, and ranked by maximum −log10(pvalue).

#### RNA Fluorescence In Situ Hybridization

RNA fluorescence *in situ* hybridization (RNA FISH) was performed on fixed brain slices with a probe to detect LepR and Pdyn RNA (RNAscope Multiplex Fluorescent Reagent Kit v2 Assay, ACD). All procedures were carried out according to the manufacturer’s instructions. Briefly, sections were pretreated with RNAscope hydrogen peroxide to block the activity of endogenous peroxidases. After a wash in distilled water, sections were permeabilized with RNAscope protease IV for 30 min at 40°C. Sections were hybridized with the Lepr and Pdyn probe at 40°C for 2 h, followed by amplification incubation steps: Amp 1, 30 min at 40°C; Amp 2, 30 min at 40°C; Amp 3, 15 min at 40°C. HRP signals were developed with RNAscope Multiplex FL v2 HRP and TSA Plus fluorophores (HRP-C1 and 1:750 TSA Plus Cy3 for Lepr, Cy2 or Cy5 for Pdyn, Cy5 for Glra2). Sections were washed with the provided washing buffer 2 × 2 min in between each step. Sections were then coverslipped with DAPI Fluoromount-G (Southern Biotech). Confocal microscope imaging was performed on a Zeiss LSM 800 microscope (Carl Zeiss).

#### Automated quantification for RNA FISH images

RNA FISH labeled cells were counted using CellProfiler image analysis software, with an analysis pipeline modified from previously published work ^106^. In brief, DAPI staining of nuclei was used to identify cells, and then cells with more than three stained speckles, or >60% of cell area covered by staining, were considered as positive for the marker.

### Behavioral assays (Figs 3-7, Extended data Figs 3-7)

#### Mice

All experiments were carried out in compliance with the Association for Assessment of Laboratory Animal Care policies and approved by the University of Virginia Animal Care and Use Committee. Animals were housed on a 12-h light/dark cycle with food (PicoLab Rodent Diet 5053) and water *ad libitum* unless otherwise indicated. All experiments were performed on male mice 12 weeks or older unless otherwise indicated. Wild-type C57BL6/J mice, LepR-Cre (B6.129-Lepr^tm3(cre)Mgmj^/J, The Jackson Laboratory #032457, RRID:IMSR_JAX:032457) ^95^, Ai14 tdTomato reporter line (B6.Cg-*Gt(ROSA)26Sor^tm14(CAG-tdTomato)Hze^*/J, Strain #007914, RRID:IMSR_JAX:007914) ^96^, and Pdyn-Cre (B6;129S-Pdyn^tm1.1(cre)Mjkr^/LowlJ, The Jackson Laboratory #027958, RRID:IMSR_JAX:027958) ^107^ mice were used.

#### Scheduled feeding (SF)

For scheduled feeding, mice were first acclimated to single housing for 7 days, followed by acclimation to IR beam interruption chambers (Columbus Instruments, or custom built ^108^) for a minimum of 72 hours before starting the recording of locomotor activity. After at least 3 days of recording of baseline locomotor activity while the mice had *ad libitum* access to food, mice were fasted at lights off (ZT12) on day 0 of SF, along with a full cage change. Mice were weighed, and injected with either vehicle (saline), 5mg/kg leptin, or 0.3mg/kg CNO at ZT2.5. Mice were then refed 3.5 hours later at ZT6 (ZT6.5 for fiber photometry experiment in Fig 6). During the first two days, mice were fed 2g, after which they were fed 2.5g. For the experiment in which we extended the food delivery window (Extended data Fig 4B-G), the same amount of food (2g on first two days, and 2.5g on days 3-10) was given to both control and extended groups. In the extended group, food was evenly split to 4 pallets and delivered at ZT6, ZT7, ZT8, ZT9. Control group received the whole pallet of food at ZT6 as other SF experiments in this work. In the leptin treatment experiments (Fig 4,5, Extended data Figs 3,4), after 5 days, treatment groups were switched, so that mice previously given saline received leptin and vice versa for the remaining 5 days. In DREADD experiments (Fig 7, Extended data Fig 6), CNO was administered for 5 days followed by 5 days of saline before switching back to 5 days of CNO administration. FAA was quantified as the amount of locomotor activity expressed in the two-hour window prior to food delivery (ZT4-6), and normalized to 24-hour activity (Figure 1, 3), or total light phase locomotor activity (in all other figures). In case of injection at ZT2.5, the 1 hour of activity post-injection was excluded to eliminate handling induced locomotion. WT groups injected with saline or leptin were age and weight-matched. Surgery operated groups were age matched.

#### Stereotactic surgery

Animals were anesthetized with isoflurane (induction 5%, maintenance 2%–2.5%; Isothesia) and placed in a stereotaxic apparatus (KOPF). A heating pad was used for the duration of the surgery to maintain body temperature and ocular lubricant was applied to the eyes to prevent desiccation. 500 nl of AAV (AAV8-hSyn-DIO-mCherry, plasmid from Addgene #44361, virus packed at UNC Vector Core ^109^; AAV8-hSyn-DIO-hM3Dq-mCherry plasmid from Addgene #50459, virus packed at UNC Vector Core; AAV1-hSyn-DIO-GCaMP7s virus from Addgene #104491-AAV1 ^110^; AAV1-CBA-DIO-GFP-TeTx ^111^ was generously gifted by Dr. Larry Zweifel [University of Washington, Seattle, WA]) was delivered using a 10 μL syringe (Hamilton) and 26-gauge needle (Hamilton) at a flow rate of 100 nl/min driven by a microsyringe pump controller (World Precision Instruments, model Micro 4). The syringe needle was left in place for 10 min and was completely withdrawn 17 min after viral delivery. For *in vivo* calcium imaging, an optic fiber guide cannula was implanted unilaterally following viral delivery, at 0.2mm dorsal to the viral injection coordinates, and stabilized on the skull with dental cement (C&B METABOND, Parkell). For electrolytic lesions, a parylene insulated, tip-exposed 2 MΩ tungsten electrode was placed bilaterally into the SCN, and a current of 1.1 mA was applied for 11 seconds. Two weeks minimum were allowed for recovery and transgene expression after surgery. Stereotaxic coordinates relative to Bregma (George Paxinos and Keith B. J. Franklin): SCN: ML: ± 0.3 mm, AP: − 0.35 mm, DV: − 5.75 mm; DMH: ML: ± 0.3 mm, AP: − 1.8 mm, DV: − 5.45 mm. After the surgery, the animals were housed individually. All surgical procedures were performed in sterile conditions and in accordance with University of Virginia IACUC guidelines.

#### Histological analysis and imaging

For fixed tissue collection, animals were deeply anesthetized (ketamine:xylazine, 280:80 mg/kg, i.p.) and perfused intracardially with ice cold 0.01 M phosphate buffer solution (PBS) followed by fixative solution (4% paraformaldehyde (PFA) in PBS at a pH of 7.4). For testing the functionality of hM3Dq (Fig 7B), 0.3mg/kg CNO was intraperitoneal injected at 2 hours prior to perfusion and brain harvesting. After perfusion, brains were harvested and post-fixed overnight at 4°C in PFA. Fixed brains were then transferred into 30% sucrose in PBS for 24 h, and then frozen on dry ice. Frozen brains were sectioned immediately or stored in −80°C for future processing. Coronal sections (30 μm) were collected with a cryostat (Microm HM 505 E). Sections were permeabilized with 0.3% Triton X-100 in PBS (PBS-T) and blocked with 3% normal donkey serum (Jackson ImmunoResearch) in PBS-T (PBS-T DS) for 30 min at room temperature. Sections were then incubated overnight at 4°C (or otherwise indicated) in primary antibodies diluted in PBS-T DS. For visualization, sections were washed with PBS-T and incubated with appropriate secondary antibodies diluted in the blocking solution for 2 h at room temperature. Sections were washed three times with PBS and mounted using DAPI Fluoromount-G (Southern Biotech). Images were captured on a Zeiss Axioplan 2 Imaging microscope equipped with an AxioCam MRm camera using AxioVision 4.6 software (Zeiss). The following primary antibodies was used for fluorescent labeling: anti-c-Fos (rabbit, 1:1k, synaptic systems #226003). The secondary antibodies (Jackson ImmunoResearch) used was Cy2-conjugated donkey anti-rabbit (1:250)(cat# 711-225-152).

#### Retrograde tracing

Rabies virus tracing: 200nl AAV1-synP-FLEX-splitTVA-EGFP-B19G (Addgene #52473-AAV1) was injected to SCN of NMS-Cre mice using the stereotactic surgery method described above. Three weeks after AAV injection, 120nl EnvA-dG-Rabies-H2B-mCherry (Salk Viral Vector Core) was delivered to the same coordinates. After one more week, fresh brains were harvested and processed for antibody labeling or RNA FISH probing as described above.

#### Bioluminescence

To determine the treatment effect on the phase of molecular circadian rhythm of the liver or SCN, PER2LUC or PER2LUC;LepR-Cre mice were individually housed under 12:12 LD light cycle for 4 consecutive days and received one of the following treatments: (1) 5mg/kg leptin at ZT6; (2) 0.3mg/kg CNO at ZT6; (3) SF feeding at ZT6; (4) saline at ZT6 or no treatment, pooled as control (used for both Fig 1B and Fig 7L). On the 5th day, mice were sacrificed between ZT5-ZT10. Brains were immediately extracted and dropped into ice cold Hanks’ Balanced Salt Solution (HBSS). After 2 minutes, brains were embedded in low melting point agarose (Precisionary Instruments, Natick, MA) and sectioned at 300 µm on a Compresstome VF-200 Vibrating Microtome (Precisionary Instruments, Natick, MA, USA). Brain region containing the SCN and liver tissue were dissected for bioluminescence recording using a method adapted from previous work ^112^. SCN slices and liver tissues were cultured in 35 mm culture dishes with 1.2 ml of DMEM (D5030, Sigma) supplemented with 3.5 g/L D-glucose, 2 mM Glutamax (Gibco), 10 mM HEPES, 25 U/ml penicillin/streptomycin, 2% B-27 Plus (Gibco), and 0.1 mM D-Luciferin sodium salt (Tocris). The culture dishes were covered with 40 mm diameter glass cover slides and sealed by high-vacuum grease (Dow Corning), and maintained in a non-humidified incubator at 36.8 °C. Bioluminescence from firefly luciferase in each of SCN slices or liver sections was recorded in 10 min intervals by a 32-channel/4-photomultiplier tube luminometer LumiCycle (Actimetrics) in the incubator. The bioluminescence data were collected and analyzed by LumiCycle Analysis software (Actimetrics). The time of the first peak in bioluminescence was manually determined from each tissue sample. Each data point represents the average of two tissue samples from the same animal which were then used for further statistical analysis.

#### In vivo fiber photometry recording

The viral vector and fiber optic implant were delivered to the target brain area within one surgical procedure as described above. Experiments were initiated at least 2 weeks post surgery to allow for recovery and transgene expression. For long-term recordings, mice were individually housed in their home cages and transferred to the recording room 3 days before experiment to acclimate to the environment. Implanted fiber-optic cannula (Thorlabs, 0.39 NA, Ø200 µm Core) was connected to fiber-optic cable (Doric Lenses, 0.37 NA, Ø200 µm Core) with the use of a zirconia mating sleeve (Doric Lenses). The fiber-optic cables were wrapped with metal sheath (Doric Lenses) to prevent breakage and connected to rotary joints (Doric Lenses) to allow free-movement. The fiber photometry system used in this work records the fluorescent signal from both calcium-dependent (465nm) and calcium-independent isosbestic (405nm) excitation light wavelengths, in which the isosbestic wavelength excitation signal is used to control for the artifacts from animal movement, fluorescent reporter expression and photobleaching. The signal was collected at 120Hz sampling rate. To limit the photobleaching of the fluorescent reporter during the long-term recording, the signal was only collected for 2 minutes every 12 minutes (For three out of seven saline treated and one out of eight leptin treated mice in the SF experiment, the data were collected for 10 minutes per hour. These datasets were included in analysis when possible. See below for details). Animals that did not exhibit Ca^2+^ signal increase in response to food presentation were considered mis-targeted and were excluded from the analysis (Extended data Fig 5C-F).

#### Fiber photometry data analysis

Data was processed in Matlab using a method modified from work described previously ^62,63^. To eliminate the artifact from motion or protein expression, the calcium-independent isosbestic (405nm excitation) signal was fitted to the calcium-dependent (465nm excitation) signal with a linear least-squares fit. The fluorescence change over baseline fluorescence (ΔF/F_0_) was calculated as (465nm induced signal - fitted 405nm induced signal) / (median of 465nm induced signal). Notably, in the long-term experiments where multiple short-term sessions were recorded, F_0_ was the median for the entire experiment for consistency. The fitted 405nm induced signal was defined as the estimated baseline, so that the values on ΔF/F_0_ curves below 0 were cutoff. For each recording session, phasic calcium signal is the integral of ΔF/F_0_ adjusted curve (Fig 6B) and represents the intracellular calcium activity during that session. The calculated phasic calcium signal was further averaged by hour and normalized to the average of the 12-hour dark phase signal in each given day. For food response during the scheduled feeding experiment, the food was delivered 1 minute after the initiation of the 2-minute recording session. The variability caused by signal strength was normalized by calculating z-score ((signal-average of baseline signal) / standard deviation of baseline signal) for food response experiments where the first 15 seconds of the 2-minute sessions were used as baseline.

#### Circadian Behavioral Analysis

Locomotor activity data were collected by wheel-equipped cages (Nalgene), or IR beam interruption chambers (Columbus Instruments, or custom built ^108^) in light-tight compartments under a 12h:12h LD cycle. Fluorescent lights (~400 lux) were used for behavioral experiments. Food and water were provided *ad libitum* unless otherwise indicated. Wheel running activity was monitored and analyzed with the ClockLab collection and analysis system (Actimetrics, Wilmette, IL). IR beam interruption activity data were analyzed in Excel for quantification or converted to a ClockLab supported file format for circadian analysis.

#### Statistical Analysis

To compare the effects of treatment over time, two-way ANOVA test was used. In experiments with a single variable and more than two groups, one-way ANOVA was performed. In case data points were missing because of technical failure (i.e. power outage during long-term recording), where repeated-measures ANOVA was not possible to perform, mixed-effects (REML) analysis was used instead. Following a significant effect in the ANOVA test, Bonferroni’s post hoc comparison was used to determine differences between individual data points. Analyses were conducted using GraphPad Prism 8 statistical software for Windows. All data were presented as means ± standard error of the mean (SEM) with p < 0.05 considered statistically significant.

## Supplemental figures

**Supplemental figure 1-.**
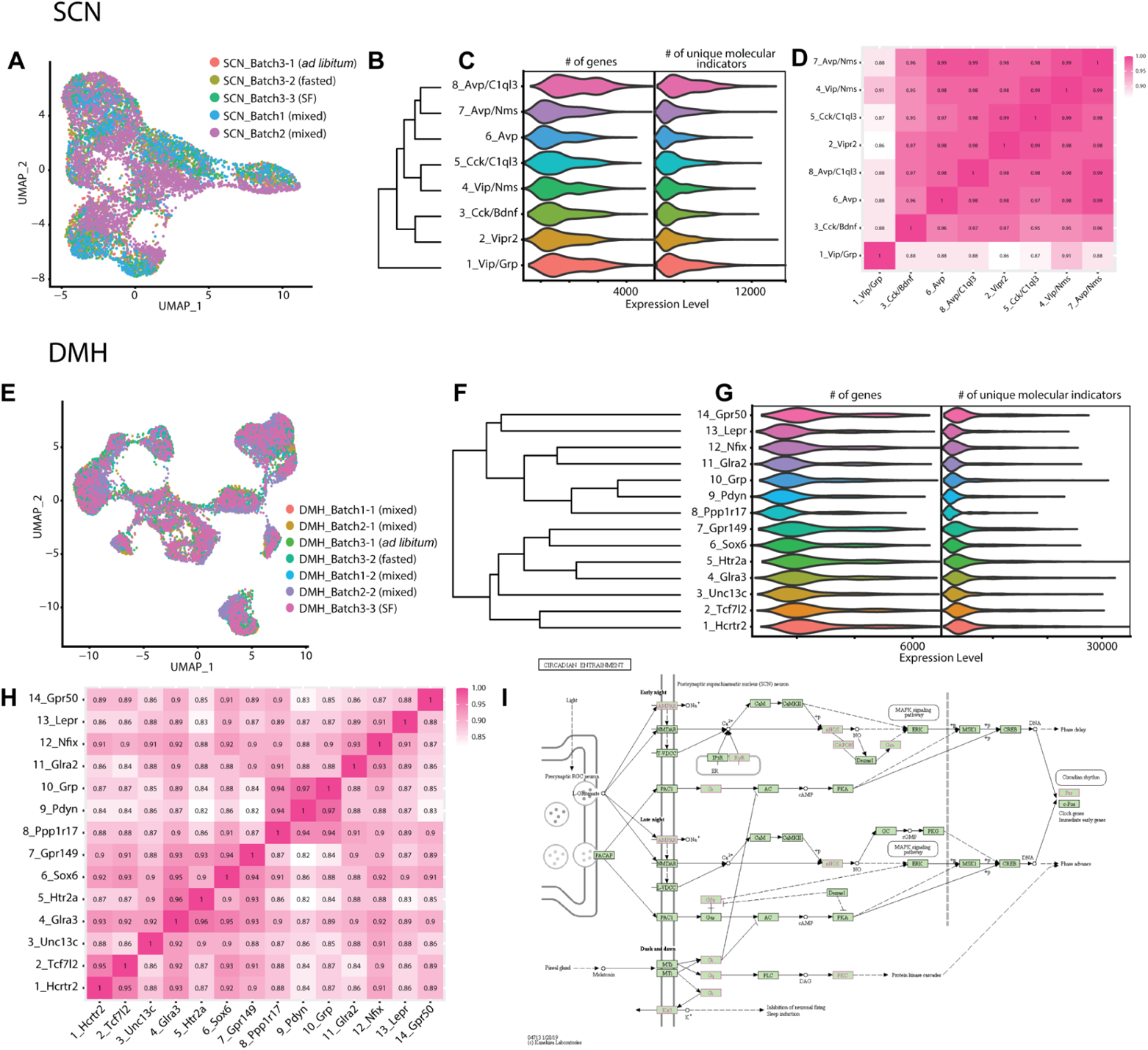
SCN and DMH single nuclei RNA-seq quality metrics and marker expression. Related to figure 1. **A.** UMAP of SCN neurons, colored by sequencing library identity, after batch correction. **B.** Phylogenetic tree indicating the relatedness of 8 SCN neuronal clusters. **C.** Expression level distribution of the number of genes per cluster (left) and number of UMIs (unique molecular identifiers, which represent unique gene transcripts; right) among SCN clusters. **D.** Correlation matrix of average expression of all genes between SCN neuronal clusters. Values within the boxes are Pearson correlation coefficients. **E.** UMAP of DMH neurons, colored by sequencing library identity, after batch correction. **F.** Phylogenetic tree indicating the relatedness of 14 DMH neuronal clusters. **G.** Expression level distribution of the number of genes per cluster (left) and number of UMIs (right) among DMH clusters. **H.** Correlation matrix of average expression of all genes within DMH neuronal clusters. Values within the boxes are Pearson correlation coefficients. **I.** KEGG “circadian entrainment” pathway map. The genes upregulated (Supplemental Fig 2D) in DMH^LepR^ neurons during SF are labeled as pink on the map.

**Supplemental figure 2-.**
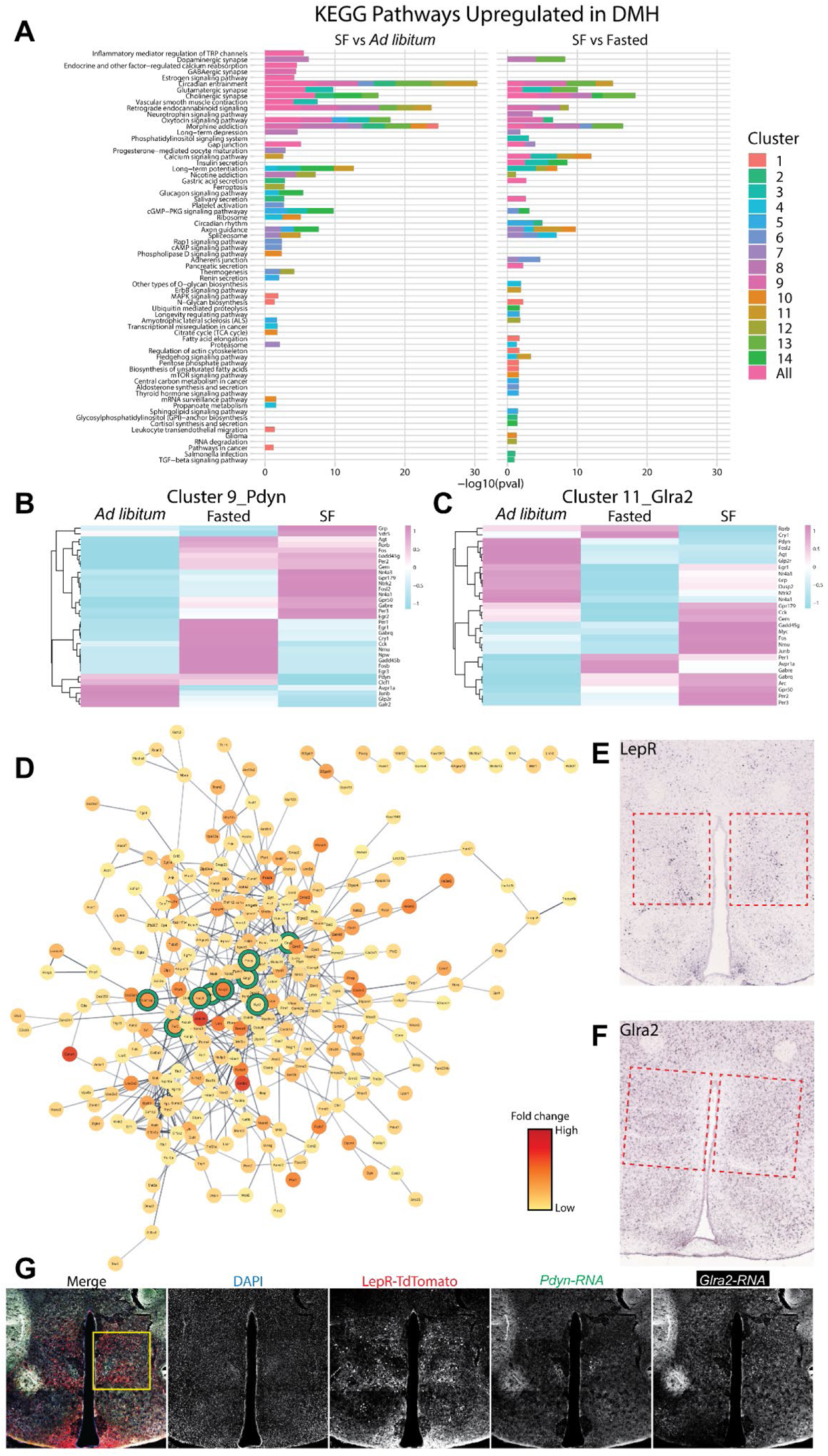
Network analysis of differentially expressed genes in the DMH between fasted and *ad libitum* conditions compared to scheduled feeding. Related to figure 2. **A.** Kyoto Encyclopedia of Genes and Genomes (KEGG) from the Mouse 2019 database comparing pathways upregulated among SF vs *ad libitum* and SF vs fasted feeding conditions in each DMH neuronal cluster and upregulated pathways in all DMH neurons. Bars are colored by clusters. Inclusion criteria required p-value <0.05 and log2 fold change >0.25. **B.** Heatmap indicating expression level of genes up and downregulated among feeding conditions in the 9_Pdyn cluster. **C.** Heatmap indicating expression level of genes up and downregulated among feeding conditions in the 11_Glra2 cluster. **D.** STRING known and predicted protein interactions using Cytoscape platform to visualize molecular networks. Input genes were derived from cluster 13_Lepr differential testing using the Wilcoxon Rank Sum Test to compare SF to *ad libitum* and SF to fasting. Inclusion criteria required False Discovery Rate <0.05 and log2 fold change >0.25. The color of circles indicates the level of average log2 fold change between SF and fasting or *ad libitum* conditions. Functional enrichment of genes involved in circadian entrainment defined by KEGG 2019 database are circled in green. **E-F.** *In situ* hybridization (ISH) of the RNA for (E) LepR, and (F) Glra2 on the brain sections that contain DMH (outlined by red boxes). The images are from Allen Mouse Brain Atlas (https://mouse.brain-map.org/search/index). **G.** Representative coronal section image showing the localization of LepR, Pdyn and Glra2 cells in the DMH. LepR cells were marked by LepR-Cre; TdTomato protein, Pdyn and Glra2 cells were marked by RNA FISH. The area indicated by the yellow box was re-imaged and presented as a zoomed-in representative image in Fig 2H.

**Supplemental figure 3-.**
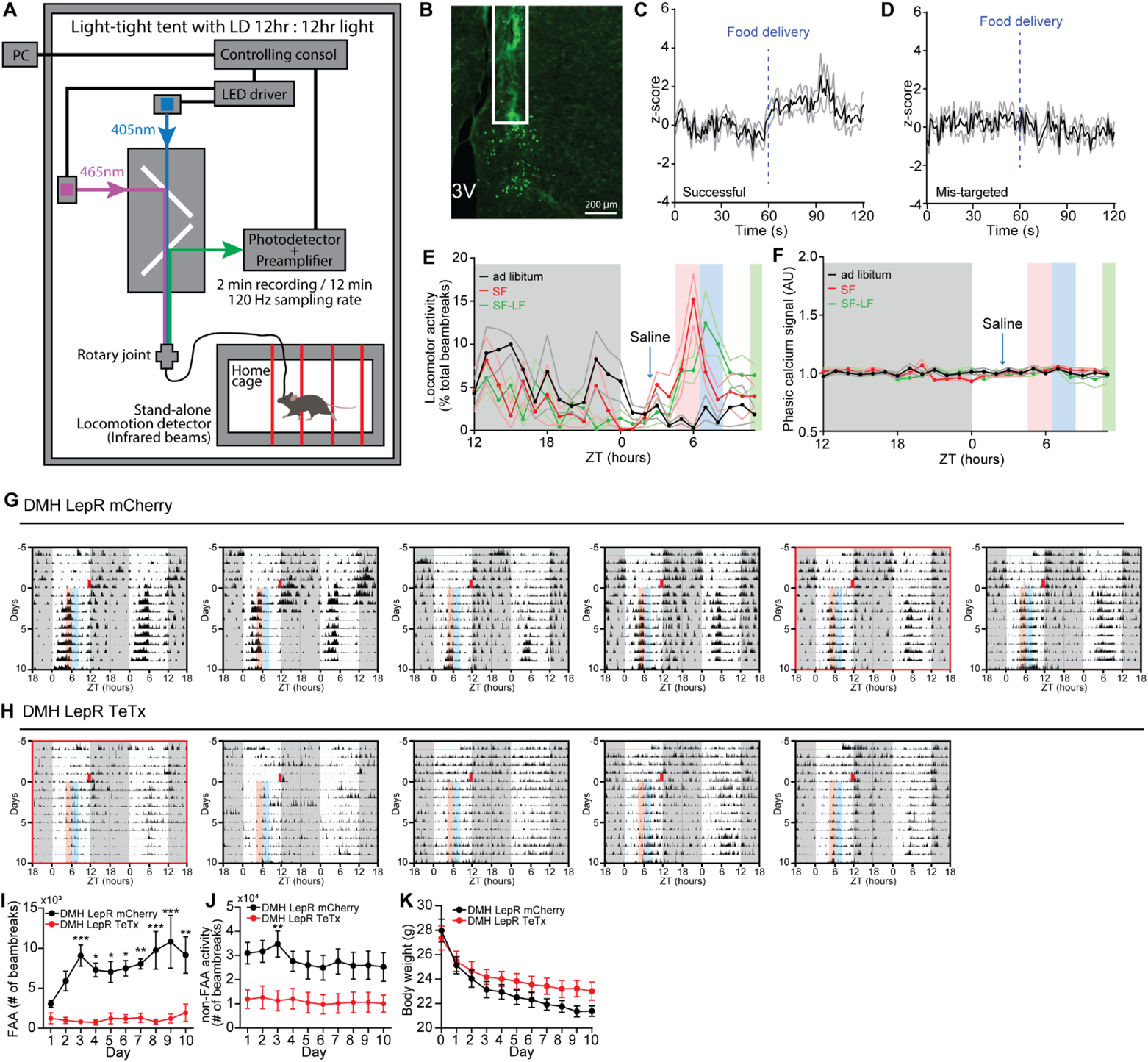
DMH^LepR^ neuron calcium recording, and TeTx silencing experiments during SF. Related to figure 3 and 4. **A.** Schematic diagram illustrating long-term fiber photometry recording coupled with locomotor activity monitoring. **B.** Representative image showing the expression of GCaMP7s in DMH^LepR^ neurons and fiber optic cannula implant (white box). **C-D.** Acute calcium signal response to food delivery on the 5th day of SF, in successful (C) and mis-targeted (D) mice. The mis-targeted animals in (D) were excluded for the analysis in Fig 3. n=6 mice/group. **E.** Locomotor activity of mis-targeted mice during SF. These mice developed strong FAA. n=5. **F.** Average phasic GCaMP7s signal of DMH^LepR^ neurons in the mis-targeted mice at 2 days before SF (black, *ad libitum*), 5th day of saline treated SF (red, SF), or 6th day of SF where saline injection was withheld and food delivery was delayed for 3.5 hours (green, SF-LF). n=6. The unaltered phasic calcium signal of these mis-targeted mice during SF suggests that the pre-meal elevation of phasic calcium signal is not due to artifact of increased locomotor activity. **G-H.** Actograms of all DMH^LepR^ mCherry (G) and DMH^LepR^ TeTx (H) mice on SF. The actograms of animals that have minimum sum of the square of the residuals to the average FAA value were used as representative figures in Fig 4, and depicted with red boxes. **I.** Absolute locomotor activity during the FAA window on SF. Repeated measures two-way ANOVA with Bonferroni post hoc comparison; n = 5-6 / group; F_virus_ (1, 9) = 41.06, p=0.0001. **J.** Absolute locomotor activity for the 22 hours per day other than FAA. Repeated measures two-way ANOVA; n = 5-6 / group; F_virus_ (1, 9) = 7.220, p=0.0249. **K.** Body weight of animals during SF. Repeated measures two-way ANOVA; n = 5-6 / group; F_virus_ (1, 9) = 1.330, p=0.2785; F_virus*time_ (10, 90) = 5.752, p<0.0001. Data are represented as mean ± SEM. *p < 0.05; **p < 0.01; ***p < 0.001; ns, not significant.

**Supplemental figure 4-.**
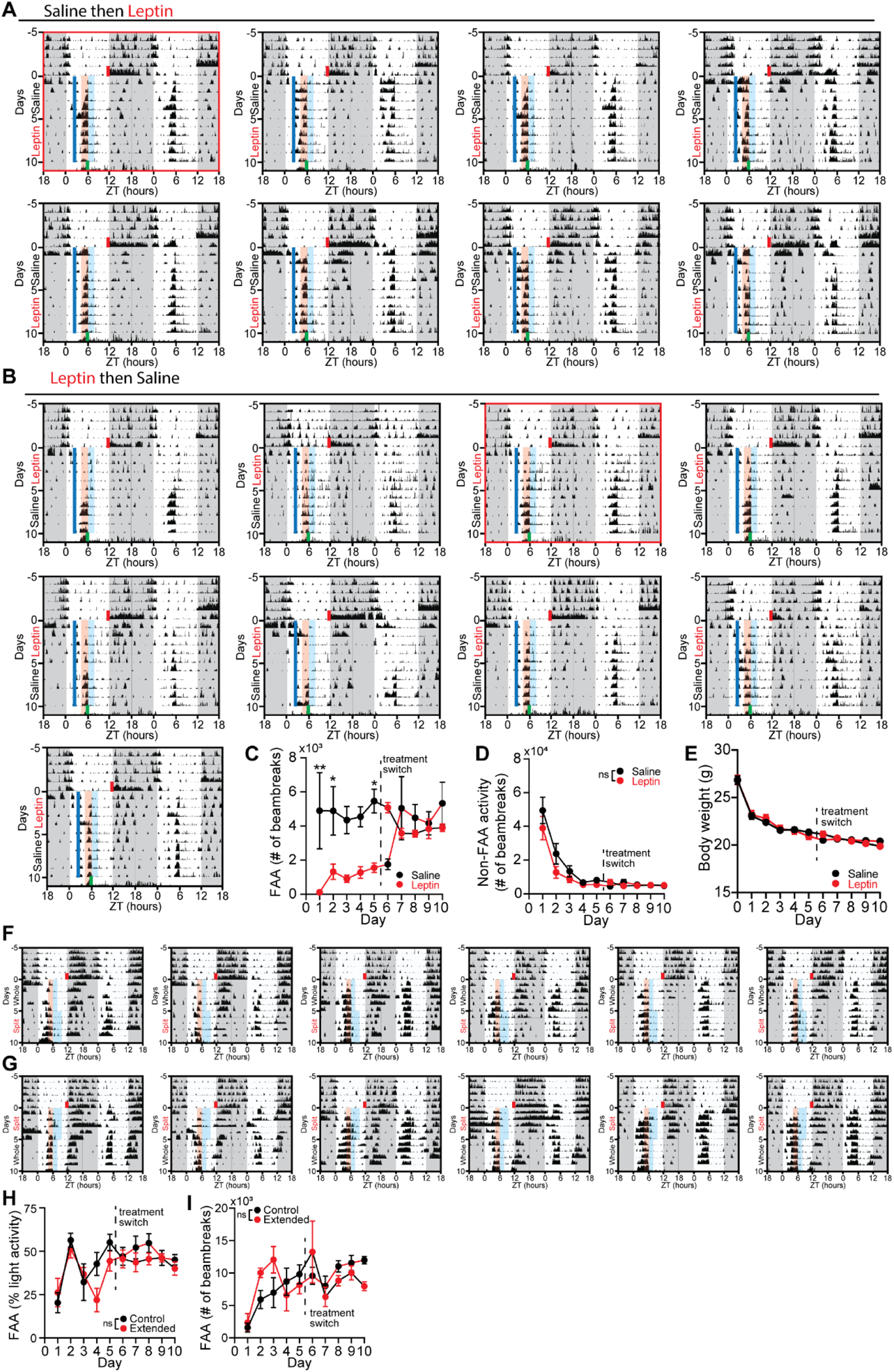
Mis-timed leptin suppresses the development but not maintenance of FAA. Related to figure 5. **A-B.** Actograms of all animals that received (A) saline then leptin or (B) leptin then saline, during scheduled feeding (SF). The actograms of animals that have minimum sum of the square of the residuals to the average FAA value were used as representative figures in Fig 5, and depicted with red boxes. **C.** Absolute locomotor activity during the FAA window, 2 hours prior to food delivery time. Repeated measures two-way ANOVA with Bonferroni post hoc comparison; n = 8-9 / group; F_treatment_ (1, 15) = 22.3, p<0.001. **D.** Absolute locomotor activity for the 22 hours per day outside of the FAA window. Repeated measures two-way ANOVA; n = 8-9 / group; F_treatment_ (1, 15) = 1.639, p=0.2198. **E.** Body weight of animals during SF. Repeated measures two-way ANOVA; n = 8-9 / group; F_treatment_ (1, 15) = 0.007550, p=0.9319; F_treatment*time_ (10, 150) = 2.633, p=0.0056. **F-G.** Actograms of all animals that received (F) control (whole pellet) then extended (same size pellet split up into 4 pellets) or (G) extended then control SF paradigm. Briefly, mice in the extended SF paradigm received one quarter of a pellet of food every hour for 4 hours, while mice in the control SF paradigm received one whole pellet at the first hour (see Methods for details). **H.** Quantification of FAA in extended SF experiment. Illustrated as a percentage of light-phase activity, without excluding any light-phase activity. Repeated measures two-way ANOVA with Bonferroni post hoc comparison; n = 6 / group; F_treatment_ (1, 10) = 0.004799, p=0.9461. **I.** Absolute locomotor activity during the FAA window in extended SF experiment. FAA window is defined as 2 hours prior to food delivery time. Repeated measures two-way ANOVA with Bonferroni post hoc comparison; n = 6 / group; F_treatment_ (1, 10) = 1.362, p=0.2703. Data are represented as mean ± SEM. *p < 0.05; **p < 0.01; ***p < 0.001; ns, not significant.

**Supplemental figure 5-.**
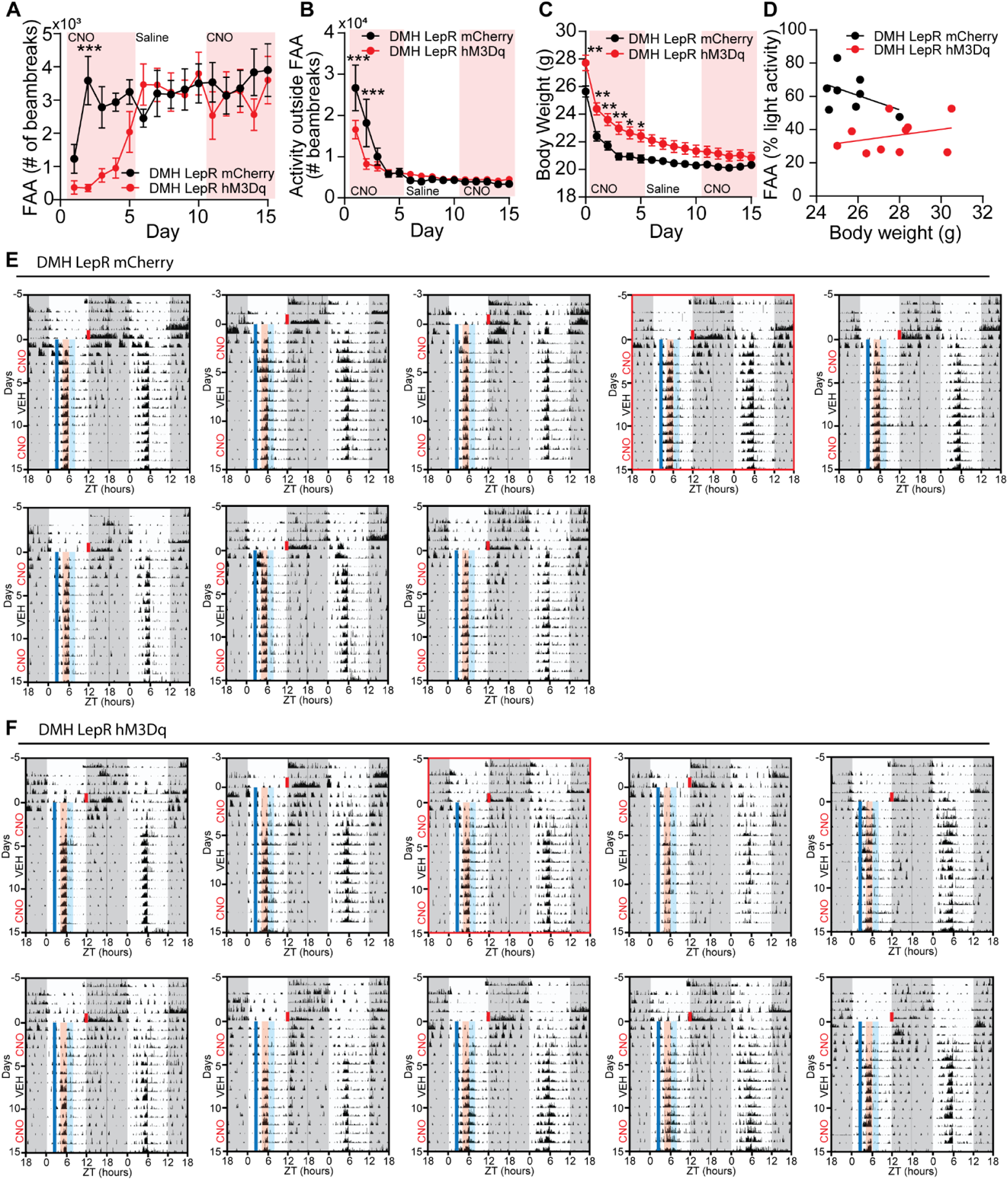
Mistimed activation of DMH^LepR^ neurons suppresses development of FAA. Related to figure 5. **A.** Absolute locomotor activity during the FAA window on SF. Repeated measures two-way ANOVA with Bonferroni post hoc comparison; n = 8-10 / group; F_time*virus_ (14, 224) = 4.647, p<0.001. **B.** Absolute locomotor activity for the 22 hours per day outside of the FAA window. Repeated measures two-way ANOVA with Bonferroni post hoc comparison; n = 8-10 / group; F_time*virus_ (14, 224) = 4.082, p<0.001. **C.** Body weight during SF. Repeated measures two-way ANOVA with Bonferroni post hoc comparison; n = 8-10 / group; F_virus_ (1, 16) = 7.342, p=0.0155. **D.** Correlation between initial body weight (day before SF) with FAA level on 5th day of SF. Higher starting body weight of hM3Dq expressing animals was not the cause of dampened development of FAA. **E-F.** Actograms of all DMH^LepR^ mCherry (E) and DMH^LepR^ hM3Dq (F) mice on SF. The actograms of animals that have minimum sum of the square of the residuals to the average FAA value were used as representative figures in Fig 5, and depicted with red boxes. Data are represented as mean ± SEM. *p < 0.05; **p < 0.01; ***p < 0.001; ns, not significant.

**Supplemental figure 6-.**
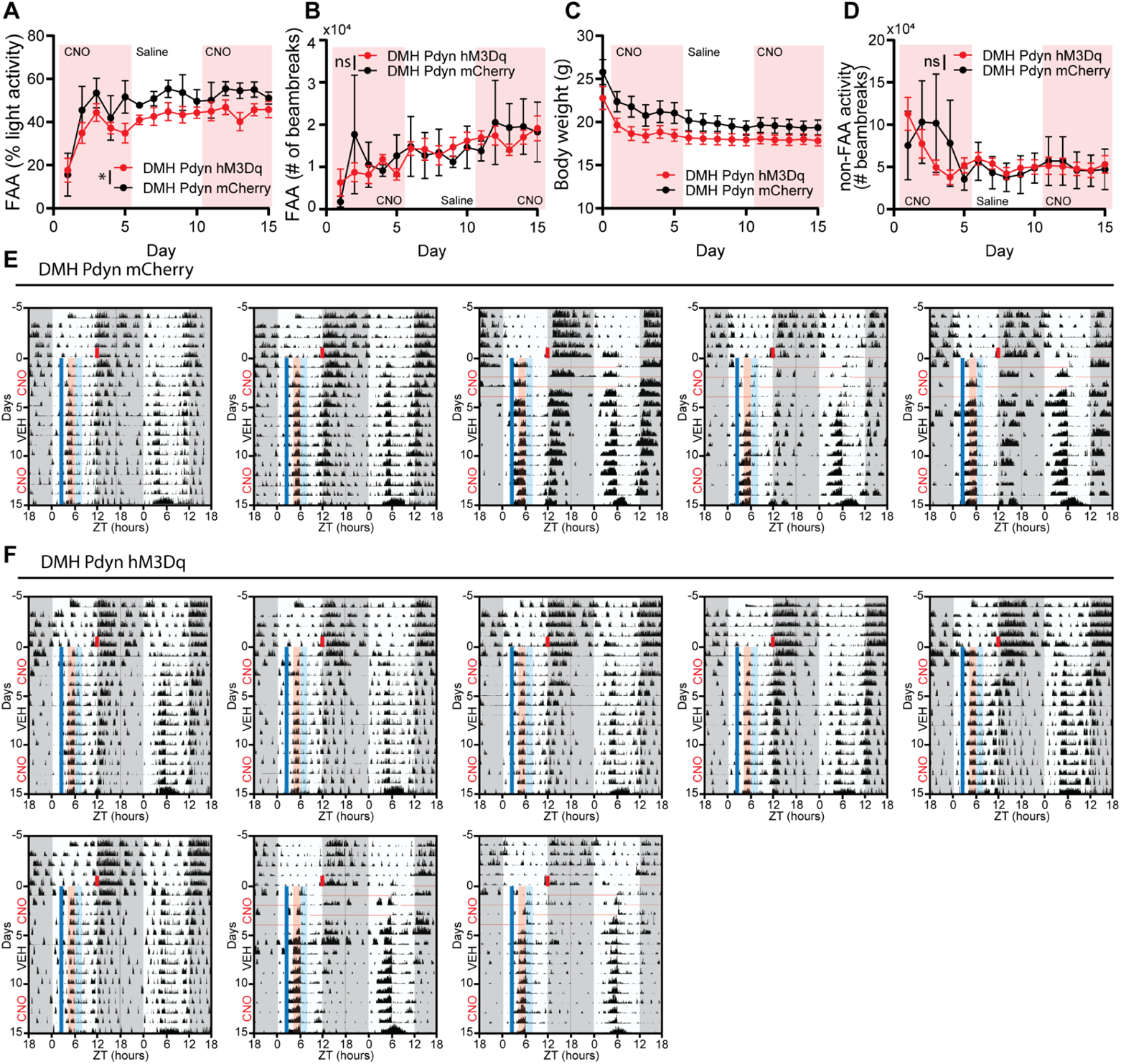
Chemogenetic activation of DMH^Pdyn^ neurons slightly inhibits the robustness of FAA. Related to figure 5. **A.** Quantification of FAA from DMH^Pdyn^ mCherry and DMH^Pdyn^ hM3Dq mice on SF that received 0.3mg/kg CNO (SF days 1-5), saline (SF days 6-10), and 0.3 mg/kg CNO (SF days 11-15) injection at ZT2.5 during SF. Pink shading indicates days with CNO injection. No shading indicates saline injection. Mixed-effects (REML) analysis with Bonferroni post hoc comparison; n = 5-8 / group; F_virus_(1, 11) = 5.601, p=0.0374. **B.** Absolute locomotor activity during the FAA window on SF. Mixed-effects (REML) analysis; n=5-8 / group; F_virus_ (1, 11) = 0.001101, p=0.9741; F_virus*time_ (14, 144) = 1.106, p=0.3576. **C.** Body weight during SF. Repeated measures two-way ANOVA with Bonferroni post hoc comparison; n = 5-8 / group; F_virus_ (1, 11) = 2.827, p=0.1208; F_time*virus_ (15, 165) = 2.834, p=0.0006. **D.** Absolute locomotor activity for the 22 hours per day outside of the FAA window. Mixed-effects (REML) analysis; n=5-8 / group; F_virus_ (1, 11) = 0.03343, p=0.8582; F_virus *time_ (14, 145) = 1.601, p=0.0853. **E-F.** Actograms of all DMH^Pdyn^ mCherry (E) and DMH^Pdyn^ hM3Dq (F) mice on SF. Horizontal red lines indicate the missing data due to technical failure. Data are represented as mean ± SEM. *p < 0.05; **p < 0.01; ***p < 0.001; ns, not significant.

**Supplemental figure 7-.**
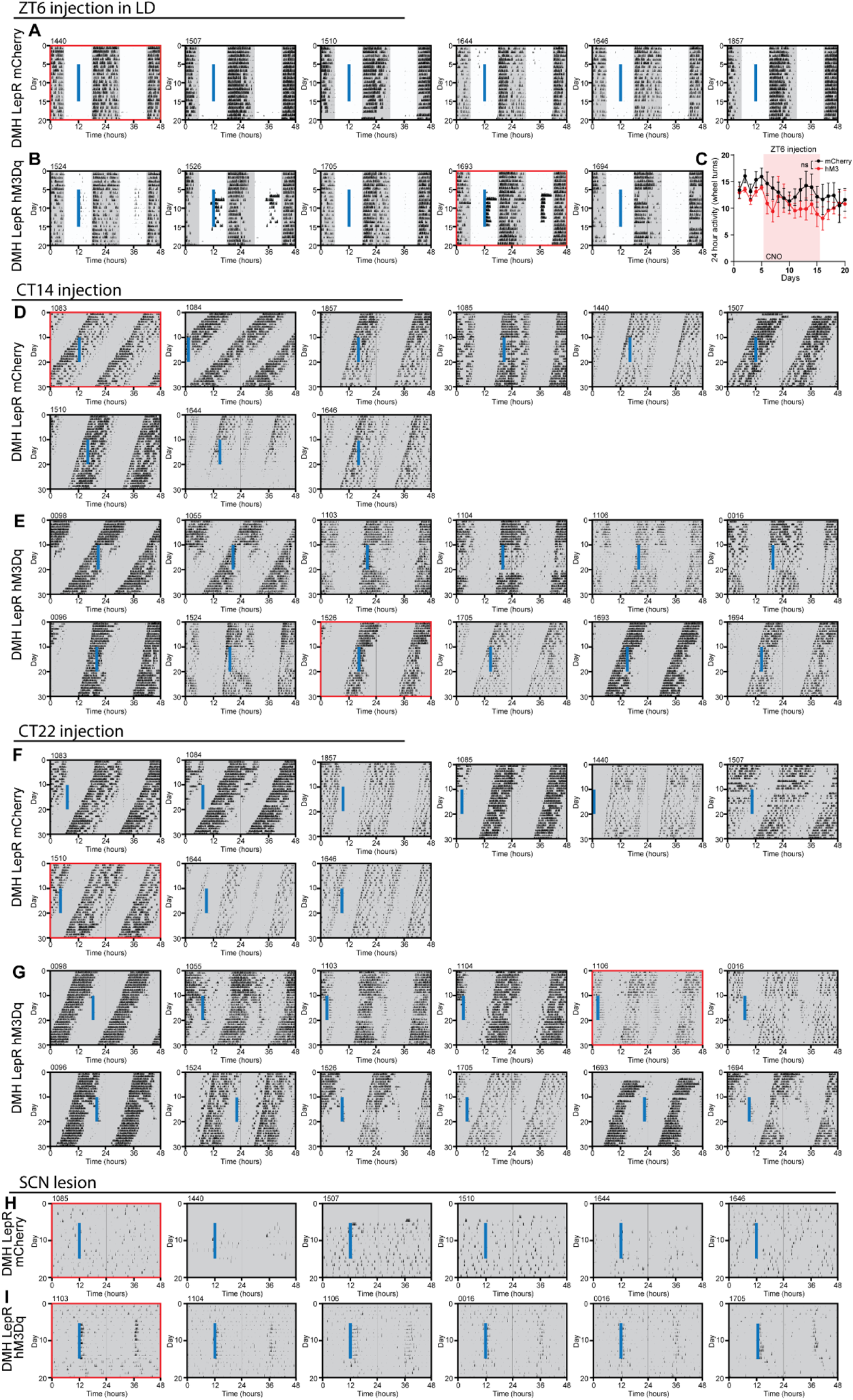
Repetitive activation of DMH^LepR^ neurons alters SCN dependent circadian locomotor activity under *ad libitum* feeding conditions. Related to figure 6 and 7. **A-B.** Actograms of all LepR Cre animals bilaterally injected with (A) AAV-hSyn-DIO-mCherry or (B) AAV-hSyn-DIO-hM3Dq-mCherry and injected with CNO at ZT6 (solid blue line) in 12-12 LD. The actograms used as representative figures in Fig 6 are depicted with red boxes. **C.** 24-hour total wheel revolutions of DMH^LepR^ mCherry and DMH^LepR^ hM3Dq animals before, during, and after CNO injections at ZT6 in 12-12 LD. Pink shading represents days with CNO injection. Repeated measures two-way ANOVA; n = 5-6 / group; F_virus_ (1, 9) = 0.7855, p=0.3985; F_virus*time_ (19, 171) = 0.9782, p=0.4890. **D-E.** Actograms of all LepR Cre animals bilaterally injected with (D) AAV-hSyn-DIO-mCherry or (E) AAV-hSyn-DIO-hM3Dq-mCherry and injected with CNO at ~CT14 (solid blue line). The actograms used as representative figures in Fig 6 are depicted with red boxes. **F-G.** Actograms of all LepR Cre animals bilaterally injected with (F) AAV-hSyn-DIO-mCherry or (G) AAV-hSyn-DIO-hM3Dq-mCherry and injected with CNO at ~CT22 (solid blue line). The actograms used as representative figures in Fig 6 are depicted with red boxes. Notably, animal #1084 in F was excluded from quantification in Fig 6N-O, because the injection time on the 10th day was near the locomotor activity onset (<3 hours) of the following day, due to the short free-running period. **H-I.** Actograms of all LepR Cre animals with electrolytic lesions of the SCN and bilaterally injected with (H) AAV-hSyn-DIO-mCherry or (I) AAV-hSyn-DIO-hM3Dq-mCherry and injected with CNO every 24 hours for 10 days (solid blue line). The actograms used as representative figures in Fig 7 are depicted with red boxes.

